# Initial assessment of the spatial learning, reversal, and sequencing task capabilities of knock- in rats with humanizing mutations in the Aβ-coding region of *App*

**DOI:** 10.1101/2022.01.24.477482

**Authors:** Hoa Pham, Tao Yin, Luciano D’Adamio

## Abstract

Model organisms mimicking the pathogenesis of human diseases are useful for identifying pathogenic mechanisms and testing therapeutic efficacy of compounds targeting them. Models of Alzheimer’s disease and related dementias aim to reproduce the brain pathology associated with these neurodegenerative disorders. Transgenic models, which involve random insertion of disease-causing genes under the control of artificial promoters, are efficient means of doing so. There are confounding factors associated with transgenic approaches, however, including target gene overexpression, dysregulation of endogenous gene expression at transgenes’ integration sites, and limitations in mimicking loss-of-function mechanisms. Furthermore, the choice of species is important, and there are anatomical, physiological, and cognitive reasons for favoring the rat over the mouse, which has been the standard for models of neurodegeneration and dementia. We report an initial assessment of the spatial learning, reversal, and sequencing task capabilities of knock-in Long-Evans rats with humanizing mutations in the Aβ-coding region of *App*, which encodes amyloid precursor protein (*App*^h/h^ rats), using the IntelliCage, an automated operant social home cage system, at 6-8 weeks of age, then again at 4-5 months of age. These rats were previously generated as control organisms for studies on neurodegeneration involving other knock-in rat models from our lab. *App*^h/h^ rats of either sex can acquire place learning and reversal tasks. They can also acquire a diagonal sequencing task by 6-8 weeks of age, but not a more advanced serial reversal task involving alternating diagonals, even by 4-5 months of age.

## Introduction

Technical innovation has enabled researchers in the past two decades to study neurodegenerative disorders with greater precision. For example, optogenetics, a technique for modulating individual neuron activity through activation of light-sensitive proteins called opsins(1), has been used to evaluate grafts of mesencephalic dopaminergic neurons derived from human embryonic stem cells in a mouse model of Parkinson’s disease(2); while electron cryo-microscopy, whose resolution has become comparable to that of X-ray crystallography(3) has revealed the structure of histopathological tau filaments in patients with frontotemporal dementia(4). Single-cell resolution transcriptomics has been used to characterize the cell diversity of the entire mouse central nervous system (CNS)(5), yielding reference data for exploring the molecular mechanisms of Alzheimer’s disease (AD), the most common form of dementia among the elderly, and related dementias in the context of animal models, which have diversified following improvements in genome-editing technologies(6). In parallel with model diversification arose high-throughput, automated methods for behavioral phenotyping(7): alongside traditional paradigms such as novel object recognition, the Morris swim task (also known as the Morris water maze), the T-maze, and the Y-maze, are operant touchscreens(8) and systems like the IntelliCage (NewBehavior AG)(9, 10), which has been used to identify cognitive deficits in multiple mouse models of AD in a socially housed setting(11).

Despite these advances and decades of research overall, the pathogenic mechanisms of AD remain poorly understood and no viable treatments have been developed(12), suggesting a fundamental flaw in the investigative approach, the underlying pathogenic assumption, or both. In this case, there is evidence implicating both: most AD research has focused on the role of amyloid-β peptide (Aβ) accumulation, either as insoluble aggregates or soluble dimers/oligomers as the causative agent of a series of events involving neuronal death, oxidative stress, synaptic loss, and neuroinflammation leading to AD, which defines the “amyloid cascade hypothesis”(13, 14). This, encouraged by links between autosomal dominant mutations in genes causing AD and changes in proteolytic processing of APP, or amyloid precursor protein(15), has prompted researchers to generate animal models of overexpression, mostly transgenic mice, as an efficient way of reproducing these alleged pathogenic conditions(16). However, individuals can have significant amyloid plaque burden without corresponding memory impairment(17–22), and many models do not exhibit the widespread neurodegeneration, cortical atrophy, or neurofibrillary tau tangles present in AD(13, 16).

Moreover, the transgenic approach is non-physiological and etiologically biased, which is not ideal for simulating the human disease state. The species also matters in evaluating the animal model paradigm. Compared to mice, rats have larger brains, which allows for more accurate direction of cannulas (for administration of drugs, biologics, viruses, etc.) and micro-dialysis probes (for sampling extracellular brain levels of neurotransmitters, Aβ, soluble tau, etc.) to individual brain regions, causing less damage and increasing specificity. In vivo brain imaging techniques, such as MRI(23) and PET(24–26), can assess the extent and course of neurodegeneration with better spatial resolution in rats. Moreover, rats are large enough for convenient in vivo electrophysiological recordings or serial sampling of cerebrospinal fluid for detection of biomarkers. Rats also express in the brain during adulthood due to alternative splicing, like humans(27), both three-(3R) and four-repeat (4R) microtubule-binding domain isoforms of tau(28–31), a protein that forms neurofibrillary tangles in AD and is mutated in frontotemporal dementia(32–39), unlike adult mice, which only express 4R isoforms(40). Finally, rats have been a choice species for behavioral, memory, and cognitive research due to their physiological similarity with humans and intelligence(41–44), which are critical when studying neurodegenerative diseases. These observations, along with the failure to translate results from current models into viable therapies(12), suggest that knock-in rat models may be better suited for the study of mechanisms underlying neurodegeneration and dementia than transgenic mouse models.

We report an assessment of the spatial learning, reversal, and sequencing task performance of knock-in Long-Evans rats carrying humanizing mutations in the Aβ-coding region of *App* (*App*^h/h^ rats) at 6-8 weeks of age, and again at 4-5 months of age, with the IntelliCage. As human and rat Aβ differ by three amino acids and Aβ aggregates are widely regarded as the main pathogenic molecules in AD with human Aβ being possibly more likely to form toxic Aβ species than rodent Aβ,(45) these animals are controls for knock-in rat models designed and generated by L. D’Adamio for neurodegeneration research, which have already yielded insights on biochemical effects of pathogenic(46) and protective(47) mutations in *APP*(48, 49), the pathogenic *PSEN1* L435F mutation(50), the Familial Danish ITM2b mutation(51), and of the R47H variant in *TREM2*(52–54), which is associated with increased AD risk (55, 56). Full characterization of these models requires cognitive and behavioral evaluation: this study establishes in part a baseline profile for the control rats while exploring analytic ideas specific to the IntelliCage paradigm that may inform further work, specifically activity curves representing aggregate behavior of experimental subjects and use of the area under those curves as a measure for statistical comparison. Moreover, the body of IntelliCage research is small: a PubMed query in December 2021 for “IntelliCage” yielded only 128 results dating from 2005, with the majority referring to mouse experiments. Therefore, this study is not only an initial cognitive characterization, but also a contribution to the nascent body of rat IntelliCage literature and a reference for analytical methods.

## Material and Methods

### Animals and Experimental Design

All experiments were done according to policies on the care and use of laboratory animals of the Ethical Guidelines for Treatment of Laboratory Animals of the NIH. Relevant protocols were approved by the Rutgers Institutional Animal Care and Use Committee (IACUC) (Protocol #201702513). All efforts were made to minimize animal suffering and reduce the number of rats used. Prior to and after behavioral analysis, males were housed 2 per cage and females were housed 3 per cage under controlled laboratory conditions with a 12-hr dark/light cycle, at a temperature of 25 ± 1°C. They were anesthetized with isoflurane, tagged subcutaneously with radio frequency identification transponders, and allowed to recover for at least a week. Rats had free access to standard rodent diet and tap water while in traditional housing and were monitored for dehydration during periods of water restriction during behavioral analysis. Long-Evans rats expressing humanized *App* alleles (*App*^h/h^) were generated as described previously.(48)

### IntelliCage

The IntelliCage for Rats (NewBehavior AG) was used to collect behavioral data. It consists of a central square home cage connected to four operant learning chambers, or *corners*. Every corner has two sides, each with a drinking bottle gated by a rotating door with a nosepoke sensor (Figure 1). The sides also include LEDs and air puff delivery valves as additional conditioning components.

**Figure 1.**
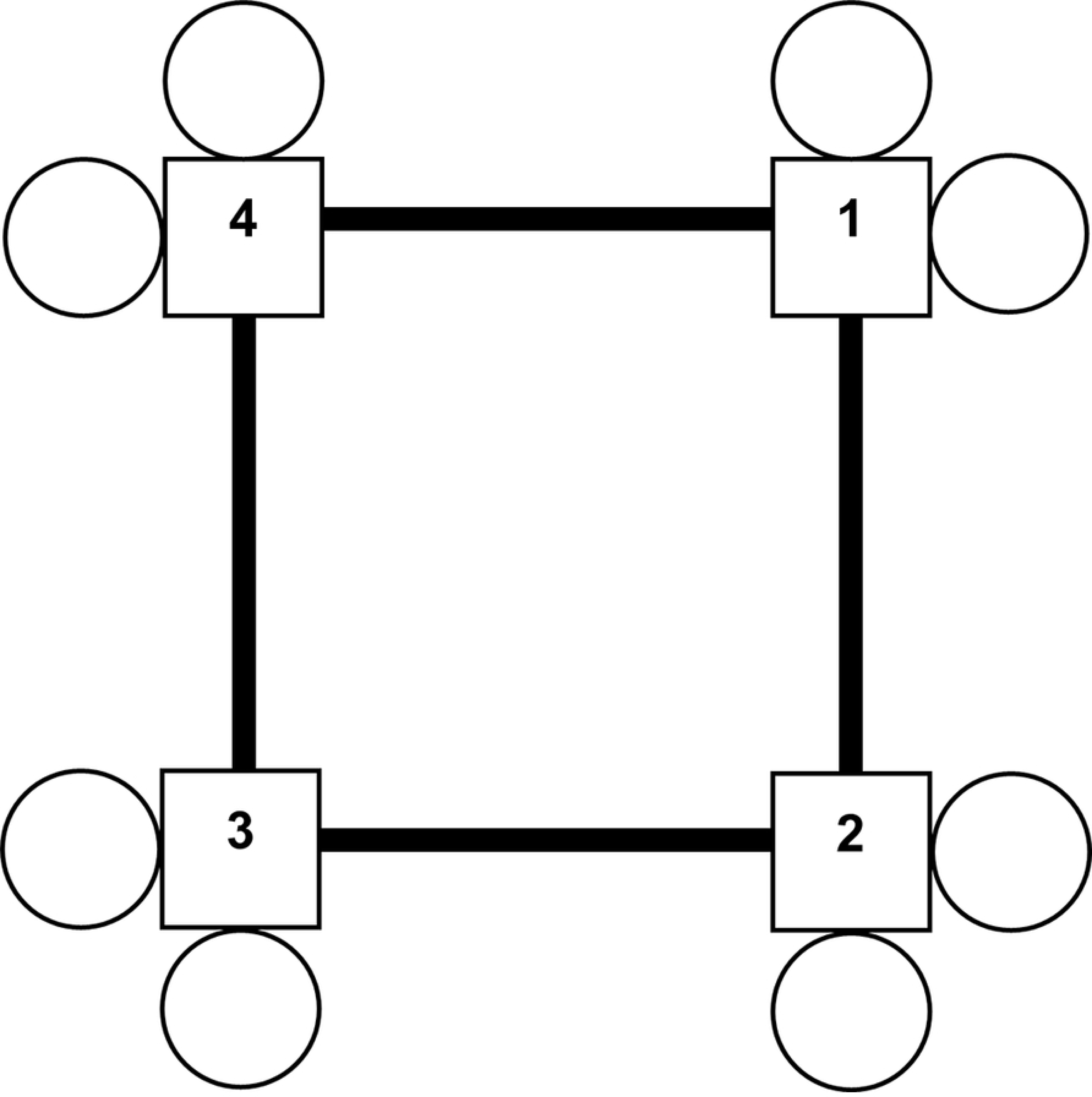
IntelliCage schematic. Central home cage and four labeled corners with two drinking sides per corner.

Behavioral programs are defined by the user within a visual coding platform. Subcutaneously injected transponders allow the IntelliCage to track the behavior of individual animals with unique radio frequency identification tags. Among the parameters tabulated for subsequent analysis are corner visits, visit lengths, visit times, number of nosepokes per visit, and number of bottle licks per visit. This system offers a variety of advantages over human observation: high-throughput, unbiased data collection and reporting; minimal risk of human error; minimal perturbation of testing conditions; and uniform testing of multiple animals simultaneously in a social setting. Two cohorts of *App*^h/h^ rats were studied, A and B, housed across four IntelliCages, separated by sex and cohort. They were run on separate program timelines, once at 6-8 weeks of age and again at 4-5 months of age (the *first pass* and *second pass* through the program timeline, respectively), as outlined in Table 1 with program descriptions in Figure 2.

**Figure 2.**
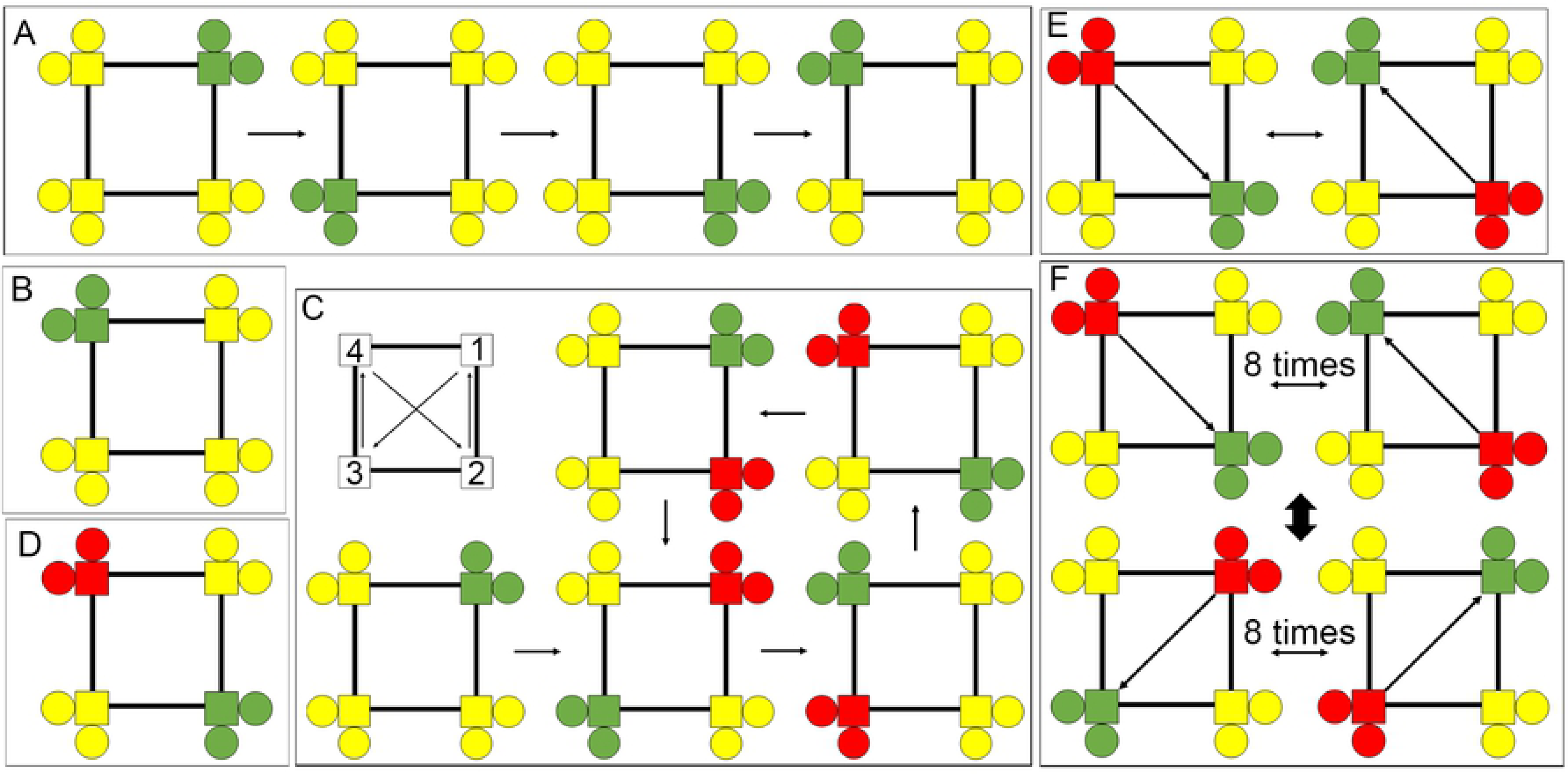
Descriptions of IntelliCage programs. (a) Single corner restriction (Cohort B only, 2 days). Progression of drinking (green) and non-drinking (yellow) corner layouts over two days. (b) Place learning (Cohort A only, 3 days). Example of correct (green) and incorrect (yellow) corner layout for a rat assigned to corner 4. (c) Place learning with corner switch (Cohort B only, 4 days). On the top left is the cycle of correct corners with movement every 45 minutes. The rest of the panel, starting from the bottom left, depicts an example of correct (green) and incorrect (yellow, or red highlighting the previously correct corner) layouts for a rat initially assigned to corner 1 and their cycle over the four phases of a drinking session, which ends with the top right layout before returning to the top center layout during the start of the next drinking session. (d) Place reversal (Cohort A only, 3 days). Example of correct (green) and incorrect (yellow, or red highlighting the previously correct corner) corner layout for a rat assigned to corner 4 during place learning. (e) Behavioral sequencing (Cohort A: 3 days, Cohort B: 5 days). Example of correct (green), lateral (yellow), and opposite (red) corner layouts and pattern for a rat initially assigned to either corner 2 or 4. (f) Serial reversal (both cohorts, 4 days). Schematic of correct (green), lateral (yellow), and opposite (red) corner layouts and pattern. The starting layout depends on the initial corner assignment.

**Table 1.**
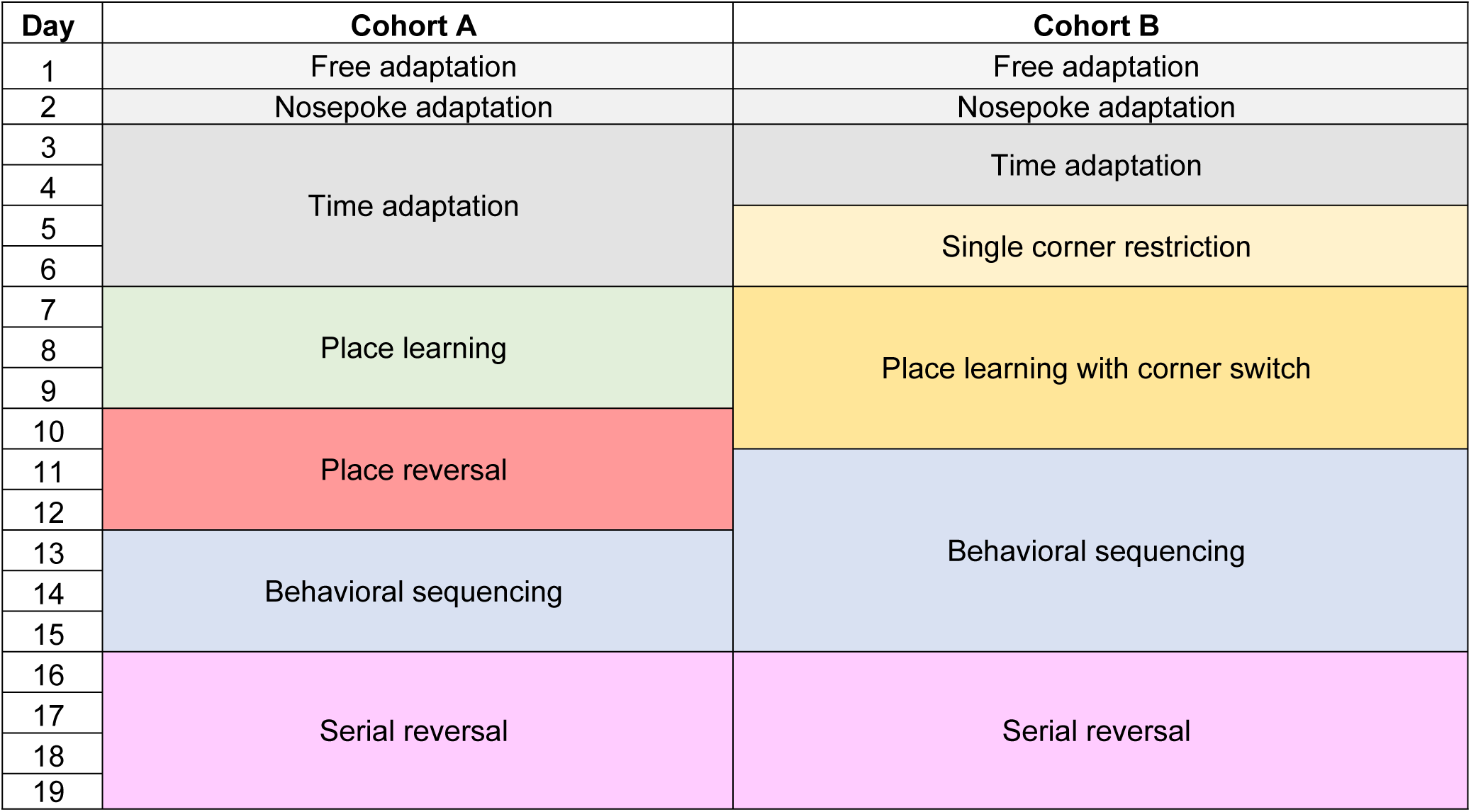
IntelliCage program timeline overview for cohorts A and B.

### IntelliCage programs

Free adaptation (both cohorts, 1 day) - The rats may drink water ad libitum and explore the IntelliCage, familiarizing themselves with its layout; all bottle access doors open in response to any corner visit.

Nosepoke adaptation (both cohorts, 1 day) - The rats learn they must activate a nosepoke sensor to open a water access door at any corner for seven seconds; this nosepoke mechanic remains active for every program hereafter.

Time adaptation (Cohort A: 4 days, Cohort B: 2 days) - The rats may only drink between 8pm and 11pm at any corner, a time window called the *drinking session*.

Single corner restriction (Cohort B only, 2 days) - All rats must drink from a single correct corner with the other corners being neutral during the drinking session. The correct corner changes after ninety minutes, such that the rats can drink at one corner during the first half of the drinking session and must switch to the opposite corner during the second half. Over two days, the correct corner designation follows the path 1->3 (1st day), then 2->4 (2nd day), covering all corners (Figure 2A).

Place learning (Cohort A only, 3 days) - The rats may only drink during the drinking session at a corner assigned to each of them; these assigned corners are considered correct, and the non- assigned corners are considered incorrect (Figure 2B).

Place learning with corner switch (Cohort B only, 4 days) - Each rat is assigned an initial correct corner where it can drink during the drinking session, as in place learning, with the other corners being incorrect. Every 45 minutes, the correct corner designations are switched according to the cycle (1->3->4->2[->1]). [Figure] illustrates this for a rat with corner 1 as the initially assigned correct corner. If corner 2 were the initial correct corner, the cycle would be shifted over once (2->1->3->4[->2]). After the first switch, the positions of the incorrect corners adjust accordingly. By the first 45-minute block of the next drinking session, the correct corner will have returned to its initial location. A *phase* refers to a 45-minute block during the drinking session in this program. The end of a phase marks when a corner switch occurs (Figure 2C).

Place reversal (Cohort A only, 3 days) - The rats may only drink during the drinking session at the corner diagonally opposite the one assigned in place learning; those reversed corners are considered correct, and the remaining corners, including the original assigned corner, are considered incorrect (Figure 2D).

Behavioral sequencing (Cohort A: 3 days, Cohort B: 5 days) - The rats must alternate between drinking at the initial learned corner and the opposite corner during the drinking session, so that one corner in the assigned diagonal is active (correct) at a time while the other is inactive (opposite); the conditions of the corners in the assigned diagonals alternate between correct and opposite, with a correct nosepoke triggering a condition switch. Visits to corners in the non-assigned diagonal are considered *lateral* visits (Figure 2E).

Serial reversal (both cohorts, 4 days) - The rats must alternate between a behavioral sequencing pattern on the original diagonal and the same on the other diagonal during the drinking session; the diagonal switches after every eight successive correct nosepokes. The corner conditions change as in the behavioral sequencing program, only now one must consider which diagonal is active in determining the corner conditions at any time (Figure 2F).

### Corner rank comparison

We followed this workflow to compare animal activity during the single corner restriction program:

1. For each 90-minute block of the program during the drinking session, rank the animals according to the number of visits made to the appropriate correct corner; there should be four lists for each IntelliCage, corresponding to the 8:00-9:30pm and 9:30-11:00pm periods of drinking sessions 1 and 2.
2. An animal is said to *out-visit* another animal at a corner if it makes more visits than the other one during a given time interval. Assign four scores to each animal equal to the number of animals it out- visits, one score for each 90-minute block; each corner should be represented once as a correct corner.
3. Use the scores to generate mean scores and standard errors for statistical analysis.

### Activity curves

We followed this workflow to produce the activity curves for each program:

1. Categorize each rat’s visits by their contextual value in the program, i.e., correct, incorrect, lateral, or opposite.
2. For each subset of rats by sex, cohort, and pass (e.g., cohort A males in their first pass), count the total number of visits those rats made for each drinking session. For a 3-day program, there should be 3 totals for a given subset.
3. For each subset as described in step 3, tabulate the fractional accumulation of visits by category over time for each drinking session, adding to each fraction, starting from 0, the value of 1 divided by the associated total count for that subset and drinking session each time a visit belonging to a certain category occurs, and 0 otherwise. For each drinking session of a given subset, the final fractions should sum to 1.
4. Match each fraction with a timestamp relative to the start of the first drinking session of a program, excluding time not belonging to a drinking session, e.g., if the *n*th fraction is associated with a visit that occurred during the 35^th^ minute of the third drinking session of a program, the fraction is matched with minute 395 (180 + 180 + 35).
5. Plot the resulting tables with time as the independent variable and fraction as the dependent variable, yielding the activity curves.

### Statistical analysis

We followed this workflow for statistical analysis of activity for each program:

1. Tabulate the fractional accumulation for individual rats as described above; in other words, make the calculations necessary to generate activity curves for each rat in the IntelliCage rather than a group of them for every drinking session.
2. Calculate the area underneath each activity curve, bounded on the left and right by the start and end of each drinking session, respectively, and below by the x-axis. Before calculating the area, we completed the curve by extending it horizontally such that the final fraction at the end of the drinking session is equal to the fraction accumulated by the last visit the animal made during that drinking session. Each result is a data point representing the cumulative activity of a specific rat for a given drinking session and visit category.
3. Run statistical tests with those data points based on desired comparisons.

## Results

### *App^h/h^* rats do not visit corners more often than other *App*^h/h^ rats during single corner restriction in cohort B at either 6-8 weeks or 4-5 months of age

After IntelliCage adaptation as outlined in Table 1, rats in cohort B were started on the single corner restriction program, which tested whether the animals were able to share this corner equally among themselves for water. The animals were assigned a rank during each 90-minute block of the two drinking sessions (four ranks total) based on visits to the actively correct corner, as described in the methods. The mean rank was used to compare animal activity (Figure 3). There were no significant differences among male rats during either pass. During the first pass one female rat (Animal 22) had a mean rank significantly lower than that of two other female rats (Animals 16 and 17), but this difference was not observed during the second pass.

**Figure 3.**
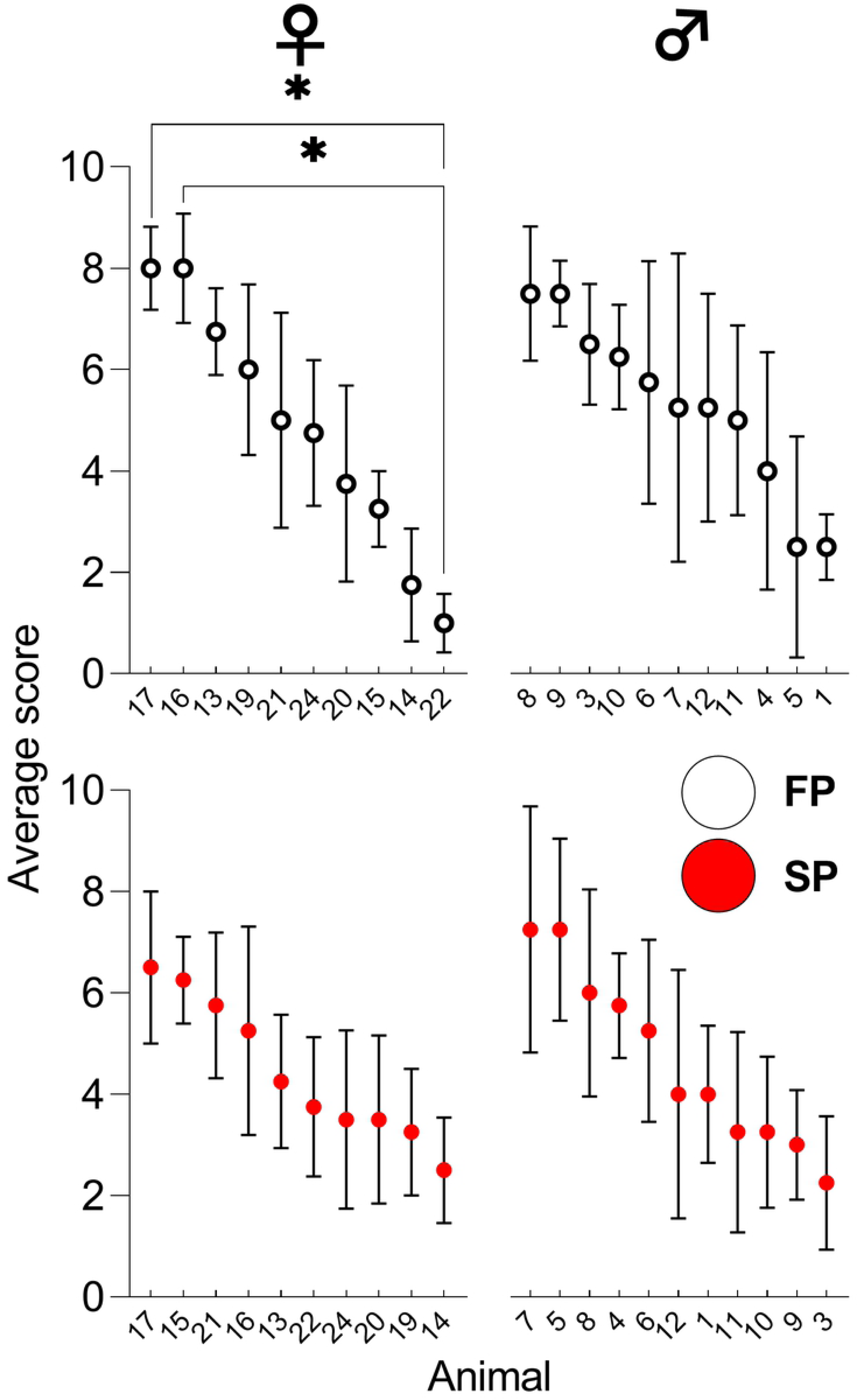
Single corner restriction, cohort B. Average scores for individual animals in cohort B by sex and pass, in decreasing order from left to right. Data points from the first and second passes are indicated by white and red circles, respectively. All data represented as mean ± SEM (\**p* < 0.05). See Table 2 for statistical analysis. FP = first pass (6-8 weeks), SP = second pass (4-5 months). ♀ = female, ♂ = male.

**Table 2.** Statistical analysis of data shown in Figure 3 for single corner restriction, cohort B. A *p*-value less than 0.05 is considered significant (\**p* < 0.05).

| Ordinary one-way ANOVA, single corner restriction, cohort B |  |  |  |
| --- | --- | --- | --- |
| Female | Pass | F (DFn, DFd) | P |
|  | First (6-8 weeks) | F (9, 30) = 3.306 | 0.0065 |
|  | Second (4-5 months) | F (9, 30) = 0.9019 | 0.5359 |
|  | post-hoc Sidak's multiple comp. test | Summary | Adjusted P |
|  | 16 (1<sup>st</sup> pass) vs. 22 (1<sup>st</sup> pass) | * | 0.0278 |
|  | 17 (1<sup>st</sup> pass) vs. 22 (1<sup>st</sup> pass) | * | 0.0278 |
| Male | Pass | F (DFn, DFd) | P |
|  | First (6-8 weeks) | F (10, 33) = 0.8408 | 0.5941 |
|  | Second (4-5 months) | F (10, 33) = 0.9633 | 0.4921 |

### *App*^h/h^ rats in cohort A can acquire a place learning task by 6-8 weeks of age with session-wise improvement

After IntelliCage adaptation as outlined in Table 1, rats in cohort A were started on the place learning program. Animal activity in this program and subsequent programs was summarized via (1) activity curves showing the fractional accumulation of corner visits by category of all animals during each drinking session, organized by sex and pass; and (2) comparisons of mean area under the activity curves of individual animals by visit category during each drinking session, to quantify differences in task performance. Qualitatively, an activity curve for correct visits that steepens as the one for incorrect visits flattens, session-wise, signifies task acquisition (Figure 4A). Analysis of area under the curves (AUC) revealed significant session-wise increases for correct visits (C-AUC) with accompanying decreases for incorrect visits (IC-AUC) for both sexes during the first pass (Figure 4B). During the second pass, there were no significant session-wise differences in C-AUC for either sex, and IC-AUC was significantly lower for the 3^rd^ drinking session compared to the 1^st^ and 2^nd^ for females alone (Figure 4B). For the 2^nd^ drinking session of the first pass, IC-AUC was significantly lower for males (Figure 4B). For females, C-AUC was significantly higher for each drinking session of the second pass compared to the first, whereas IC-AUC was significantly lower for the 1^st^ drinking session during the second pass; there were no significant differences between passes for males (Figure 4C).

**Figure 4.**
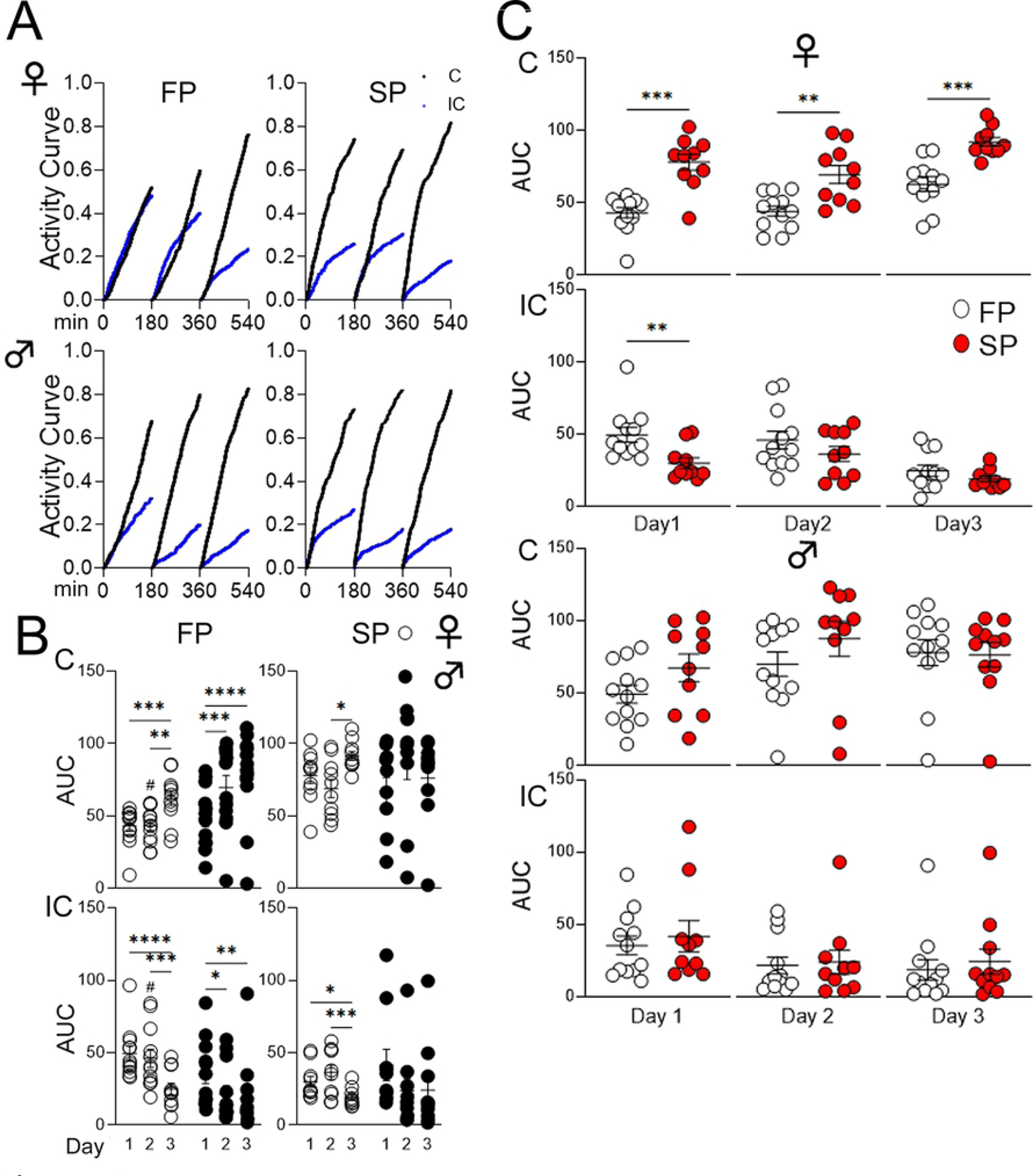
Place learning, cohort A. **(A)** Activity curves showing the fractional accumulation (y-axis) of visits over drinking session time (x-axis), reset every 180 minutes, by sex and pass. Curves for correct and incorrect visits are black and blue, respectively. **(B)** Sex comparison of area under the curve (AUC) for activity curves of individual animals by visit category and pass. “*” denotes significant comparisons within sex across program days, while “#” denotes those for the corresponding program day between sexes. Female (♀) and male (♂) data points are indicated by white and black circles, respectively. **(C)** Pass comparison of AUCs for activity curves of individual animals by sex, program day, and visit category. Data points from the first and second passes are indicated by white and red circles, respectively. All data represented as mean ± SEM (\**p* < 0.05, \*\**p* < 0.01, \*\*\**p* < 0.001, \*\*\*\**p* < 0.0001, similarly for ^#^*p* < 0.05, etc.). See Tables 3-5 for statistical analysis. FP = first pass (6-8 weeks), SP = second pass (4-5 months), C = correct (visits), IC = incorrect (visits).

**Table 3.** Statistical analysis of data shown in Figure 4B for place learning, cohort A, first pass (6-8 weeks). A *p*-value less than 0.05 is considered significant (*^/#^*p* < 0.05, \*\**p* < 0.01, \*\*\**p* < 0.001, \*\*\*\**p* < 0.0001). ns = not significant.

| Figure 4B | Mixed-effects analysis, place learning, cohort A, first pass |  |  |
| --- | --- | --- | --- |
| Correct<br>(C-AUC) | Source of Variation | F (DFn, DFd) | P |
|  | Interaction | F (2, 43) = 4.389 | 0.0184 |
|  | Day | F (2, 43) = 24.08 | <0.0001 |
|  | Sex | F (1, 22) = 3.847 | 0.0626 |
|  | post-hoc Sidak's multiple comp. test | Summary | Adjus. P |
|  | Female, day 1 vs. day 2 | ns | 0.9995 |
|  | Female, day 1 vs. day 3 | *** | 0.0010 |
|  | Female, day 2 vs. day 3 | ** | 0.0013 |
|  | Male, day 1 vs. day 2 | *** | 0.0003 |
|  | Male, day 1 vs. day 3 | **** | <0.0001 |
|  | Male, day 2 vs. day 3 | ns | 0.2879 |
| Incorrect<br>(IC-AUC) | Source of Variation | F (DFn, DFd) | P |
|  | Interaction | F (2, 43) = 3.207 | 0.0503 |
|  | Day | F (2, 43) = 17.84 | <0.0001 |
|  | Sex | F (1, 22) = 4.410 | 0.0474 |
|  | post-hoc Sidak's multiple comp. test | Summary | Adjus. P |
|  | Female, day 1 vs. day 2 | ns | 0.8453 |
|  | Female, day 1 vs. day 3 | **** | <0.0001 |
|  | Female, day 2 vs. day 3 | *** | 0.0006 |
|  | Male, day 1 vs. day 2 | * | 0.0136 |
|  | Male, day 1 vs. day 3 | ** | 0.0023 |
|  | Male, day 2 vs. day 3 | ns | 0.9004 |
|  | Female vs. Male, day 1 | ns | 0.2537 |
|  | Female vs. Male, day 2 | # | 0.0130 |
|  | Female vs. Male, day 3 | ns | 0.7167 |

**Table 4.** Statistical analysis of data shown in Figure 4B for place learning, cohort A, second pass (4-5 months). A *p*-value less than 0.05 is considered significant (\**p* < 0.05, \*\*\**p* < 0.001). ns = not significant.

| Figure 4B | Mixed-effects analysis, place learning, cohort A, second pass |  |  |
| --- | --- | --- | --- |
| Correct<br>(C-AUC) | Source of Variation | F (DFn, DFd) | P |
|  | Interaction | F (2, 35) = 4.975 | 0.0125 |
|  | Day | F (2, 35) = 2.064 | 0.1422 |
|  | Sex | F (1, 20) = 0.2437 | 0.6269 |
| Incorrect<br>(IC-AUC) | Source of Variation | F (DFn, DFd) | P |
|  | Interaction | F (2, 35) = 5.580 | 0.0079 |
|  | Day | F (2, 35) = 6.305 | 0.0046 |
|  | Sex | F (1, 20) = 0.1981 | 0.6610 |
|  | post-hoc Sidak's multiple comp. test | Summary | Adjus. P |
|  | Female, day 1 vs. day 2 | ns | 0.3829 |
|  | Female, day 1 vs. day 3 | * | 0.0470 |
|  | Female, day 2 vs. day 3 | *** | 0.0009 |
|  | Male, day 1 vs. day 2 | ns | 0.0632 |
|  | Male, day 1 vs. day 3 | ns | 0.1004 |
|  | Male, day 2 vs. day 3 | ns | 0.9945 |

**Table 5.** Statistical analysis of data shown in Figure 4C for place learning, cohort A, pass comparison. A *p*-value less than 0.05 is considered significant (\*\**p* < 0.01, \*\*\**p* < 0.001). ns = not significant.

| Figure 4C | Paired *t* tests, place learning, cohort A, pass comparison |  |  |  |
| --- | --- | --- | --- | --- |
| Correct<br>(C-AUC) | Comparison | *t*, *df* | Summary | *P* |
|  | Female, day 1 | *t*=5.672, *df*=9 | *** | 0.0003 |
|  | Male, day 1 | *t*=1.141, *df*=9 | ns | 0.2834 |
|  | Female, day 2 | *t*=4.655, *df*=9 | ** | 0.0012 |
|  | Male, day 2 | *t*=0.9056, *df*=9 | ns | 0.3888 |
|  | Female, day 3 | *t*=5.332, *df*=9 | *** | 0.0005 |
|  | Male, day 3 | *t*=0.01567, *df*=10 | ns | 0.9878 |
| Incorrect<br>(IC-AUC) | Comparison | *t*, *df* | Summary | *P* |
|  | Female, day 1 | *t*=3.283, *df*=9 | ** | 0.0095 |
|  | Male, day 1 | *t*=1.055, *df*=9 | ns | 0.3189 |
|  | Female, day 2 | *t*=0.5384, *df*=9 | ns | 0.6034 |
|  | Male, day 2 | *t*=0.4251, *df*=9 | ns | 0.6807 |
|  | Female, day 3 | *t*=0.7488, *df*=9 | ns | 0.4731 |
|  | Male, day 3 | *t*=0.3990, *df*=10 | ns | 0.6983 |

### *App*^h/h^ rats in cohort A can acquire a place reversal task by 6-8 weeks of age with session-wise improvement

After place learning, rats in cohort A were started on the place reversal program, which switches the correct corner in place learning to the one diagonally opposing it. Activity curves showed qualitative improvement, like those shown for place learning (Figures 4A and 5A). There were also significant session-wise increases in C-AUC and decreases in IC-AUC for both sexes during the first pass, with no significant differences seen during the second pass for either sex (Figure 5B).

**Figure 5.**
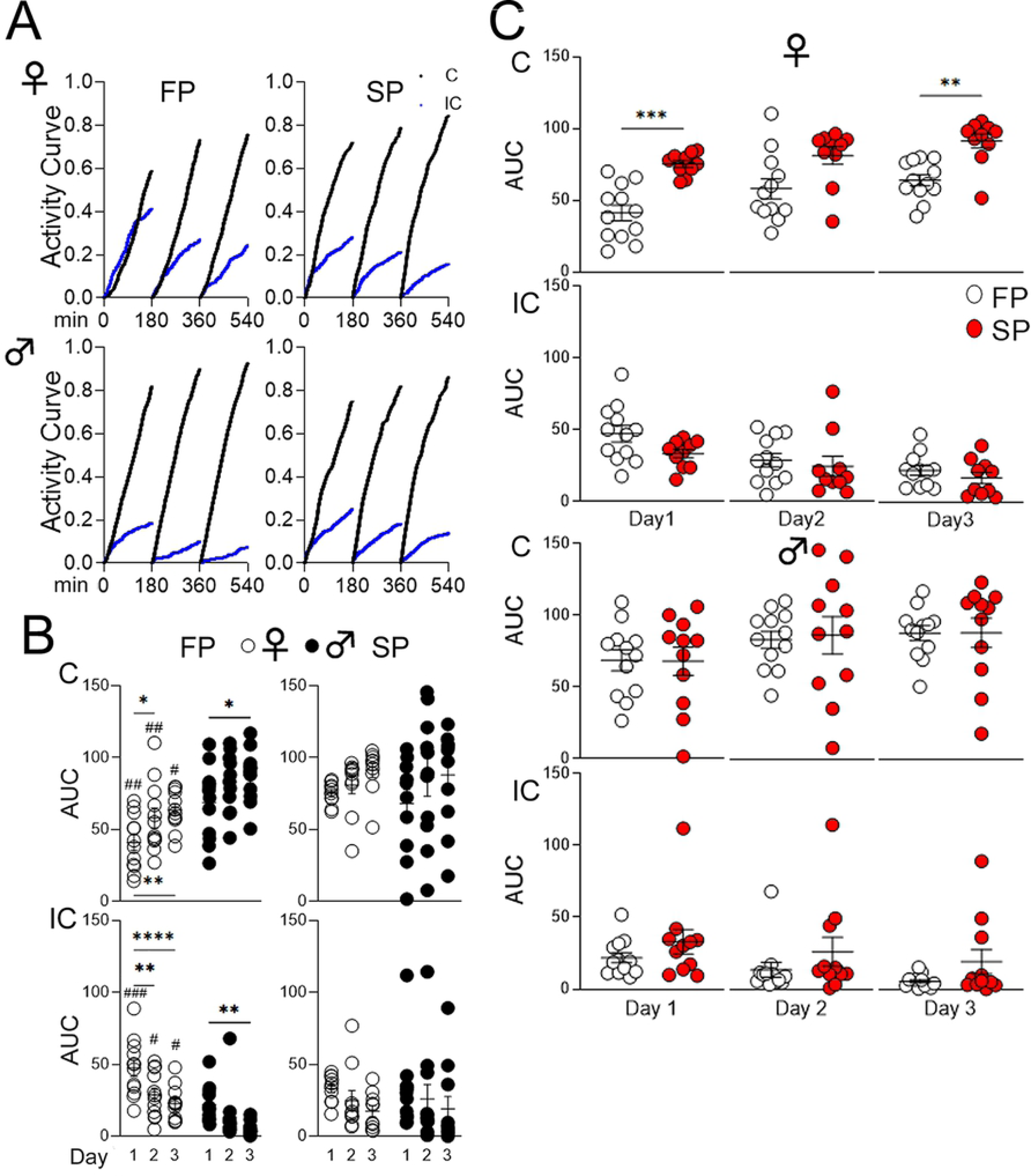
Place reversal, cohort A. **(A)** Activity curves showing the fractional accumulation (y-axis) of visits over drinking session time (x-axis), reset every 180 minutes, by sex and pass. Curves for correct and incorrect visits are black and blue, respectively. **(B)** Sex comparison of area under the curve (AUC) for activity curves of individual animals by visit category and pass. “*” denotes significant comparisons within sex across program days, while “#” denotes those for the corresponding program day between sexes. Female (♀) and male (♂) data points are indicated by white and black circles, respectively. **(C)** Pass comparison of AUCs for activity curves of individual animals by sex, program day, and visit category. Data points from the first and second passes are indicated by white and red circles, respectively. All data represented as mean ± SEM (\**p* < 0.05, \*\**p* < 0.01, \*\*\**p* < 0.001, \*\*\*\**p* < 0.0001, similarly for ^#^*p* < 0.05, etc.). See Tables 6-8 for statistical analysis. FP = first pass (6-8 weeks), SP = second pass (4-5 months), C = correct (visits), IC = incorrect (visits).

There were significant sex differences seen during the first pass, with C-AUC higher and IC-AUC lower for males for every drinking session, but not during the second pass (Figure 5B). For females, C-AUC was significantly higher during the second pass compared to the first for the 1^st^ and 3^rd^ drinking sessions, with the value for the 2^nd^ drinking session being higher but not reaching significance, while IC-AUC was not significantly different for any drinking session; there were no significant differences between passes for
males (Figure 5C).

**Table 6.** Statistical analysis of data shown in Figure 5B for place reversal, cohort A, first pass (6-8 weeks). A *p*-value less than 0.05 is considered significant (*^/#^*p* < 0.05, **^/##^*p* < 0.01, ^###^*p* < 0.001, \*\*\*\**p* < 0.0001). ns = not significant.

| Figure 5B | Two-way RM ANOVA, place reversal, cohort A, first pass |  |  |
| --- | --- | --- | --- |
| Correct<br>(C-AUC) | Source of Variation | F (DFn, DFd) | P |
|  | Interaction | F (2, 44) = 0.08855 | 0.9154 |
|  | Day | F (2, 44) = 12.26 | <0.0001 |
|  | Sex | F (1, 22) = 15.38 | 0.0007 |
|  | post-hoc Sidak's multiple comp. test | Summary | Adjusted P |
|  | Female, day 1 vs. day 2 | * | 0.0277 |
|  | Female, day 1 vs. day 3 | ** | 0.0020 |
|  | Female, day 2 vs. day 3 | ns | 0.7307 |
|  | Male, day 1 vs. day 2 | ns | 0.0713 |
|  | Male, day 1 vs. day 3 | * | 0.0109 |
|  | Male, day 2 vs. day 3 | ns | 0.8461 |
|  | Female vs. Male, day 1 | ## | 0.0042 |
|  | Female vs. Male, day 2 | ## | 0.0100 |
|  | Female vs. Male, day 3 | # | 0.0153 |
| Incorrect<br>(IC-AUC) | Source of Variation | F (DFn, DFd) | P |
|  | Interaction | F (2, 44) = 1.091 | 0.3449 |
|  | Day | F (2, 44) = 16.39 | <0.0001 |
|  | Sex | F (1, 22) = 21.79 | 0.0001 |
|  | post-hoc Sidak's multiple comp. test | Summary | Adjusted P |
|  | Female, day 1 vs. day 2 | ** | 0.0026 |
|  | Female, day 1 vs. day 3 | **** | <0.0001 |
|  | Female, day 2 vs. day 3 | ns | 0.5360 |
|  | Male, day 1 vs. day 2 | ns | 0.2942 |
|  | Male, day 1 vs. day 3 | ** | 0.0083 |
|  | Male, day 2 vs. day 3 | ns | 0.3454 |
|  | Female vs. Male, day 1 | ### | 0.0002 |
|  | Female vs. Male, day 2 | # | 0.0319 |
|  | Female vs. Male, day 3 | # | 0.0154 |

**Table 7.**
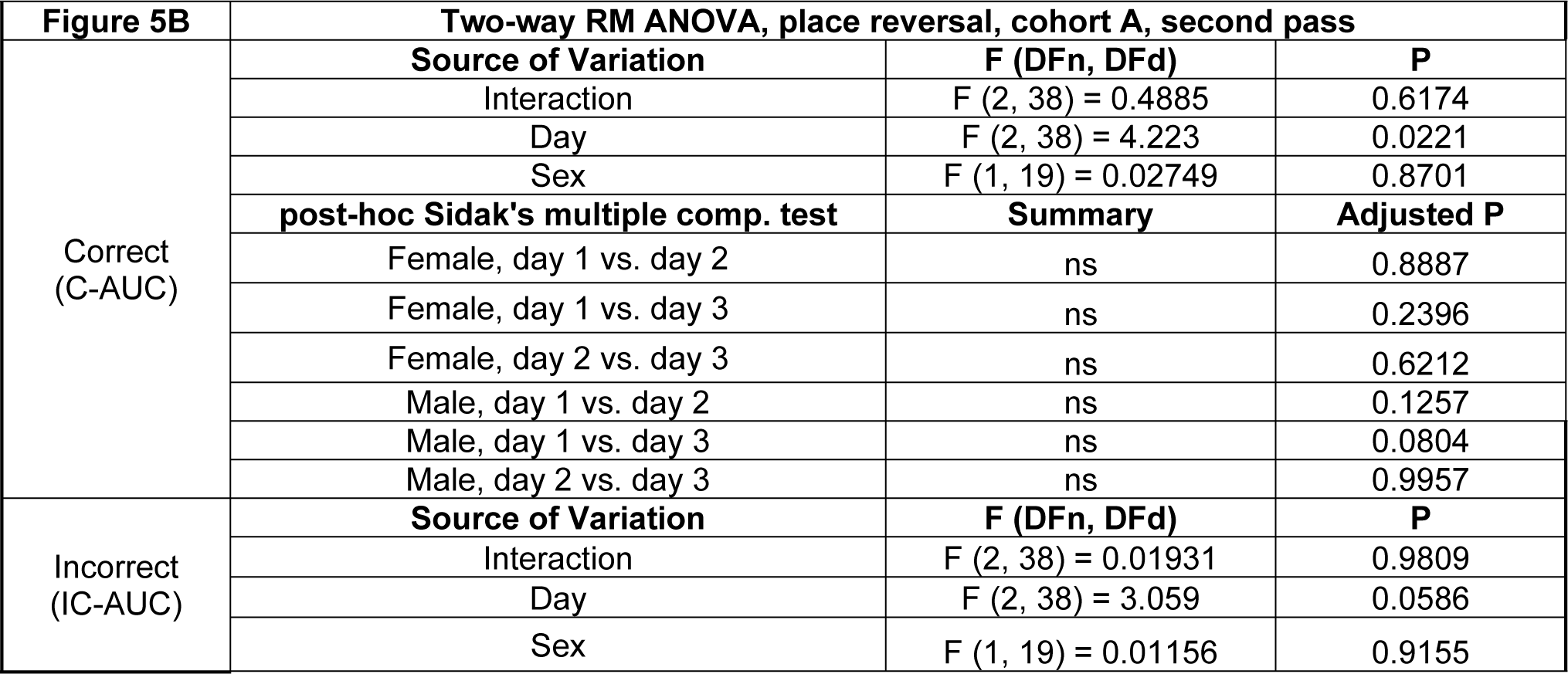
Statistical analysis of data shown in Figure 5B for place reversal, cohort A, second pass (4-5 months). A *p*-value less than 0.05 is considered significant. ns = not significant.

**Table 8.** Statistical analysis of data shown in Figure 5C for place reversal, cohort A, pass comparison. A *p*-value less than 0.05 is considered significant (\*\**p* < 0.01, \*\*\**p* < 0.001). ns = not significant.

| Figure 5C | Paired *t* tests, place reversal, cohort A, pass comparison |  |  |  |
| --- | --- | --- | --- | --- |
| Correct<br>(C-AUC) | Comparison | *t*, *df* | Summary | *P* |
|  | Female, day 1 | *t*=5.188, *df*=9 | *** | 0.0006 |
|  | Male, day 1 | *t*=0.06592, *df*=10 | ns | 0.9487 |
|  | Female, day 2 | *t*=1.595, *df*=9 | ns | 0.1453 |
|  | Male, day 2 | *t*=0.3032, *df*=10 | ns | 0.7680 |
|  | Female, day 3 | *t*=3.329, *df*=9 | ** | 0.0088 |
|  | Male, day 3 | *t*=0.09445, *df*=10 | ns | 0.9266 |
| Incorrect<br>(IC-AUC) | Comparison | *t*, *df* | Summary | *P* |
|  | Female, day 1 | *t*=1.661, *df*=9 | ns | 0.1310 |
|  | Male, day 1 | *t*=1.387, *df*=10 | ns | 0.1955 |
|  | Female, day 2 | *t*=0.003776, *df*=9 | ns | 0.9971 |
|  | Male, day 2 | *t*=1.032, *df*=10 | ns | 0.3265 |
|  | Female, day 3 | *t*=0.6898, *df*=9 | ns | 0.5077 |
|  | Male, day 3 | *t*=1.668, *df*=10 | ns | 0.1263 |

### *App*^h/h^ rats in cohort B can acquire a place learning with corner switching task by 4-5 months of age with session-wise improvement

Rather than progressing from place learning to place reversal as cohort A rats did, cohort B rats were started on the place learning with corner switch program after single corner restriction. This program was designed to be a faster-paced combination of place learning and place reversal, with correct corners switching every 45 minutes within a drinking session. Activity curves show marked differences between passes for both sexes, with curves for correct visits surpassing those for incorrect visits earlier during the second pass; notably, for females, the correct curve surpassed the incorrect curve by the end of the 2^nd^ drinking session during the second pass, whereas the correct curve never surpassed the incorrect one during the first pass (Figure 6A). No significant session-wise differences in AUC were seen during the first pass for either sex, but there were significant increases in C-AUC and decreases in IC-AUC during the second pass for both sexes, more pronounced for C-AUC (Figure 6B). Sex differences were significant for each drinking session during the second pass, with C-AUC higher for females and IC-AUC lower for males (Figure 6B). For females during the second pass compared to the first pass, C-AUC was significantly higher for females for all but the 1^st^ drinking session, while IC-AUC was only significantly different for the 4^th^ drinking session (Figure 6C). For males during the second pass compared to the first pass, C-AUC was significantly higher for all drinking sessions, while IC-AUC was significantly lower for all drinking sessions except the 2^nd^ (Figure 6C).

**Figure 6.**
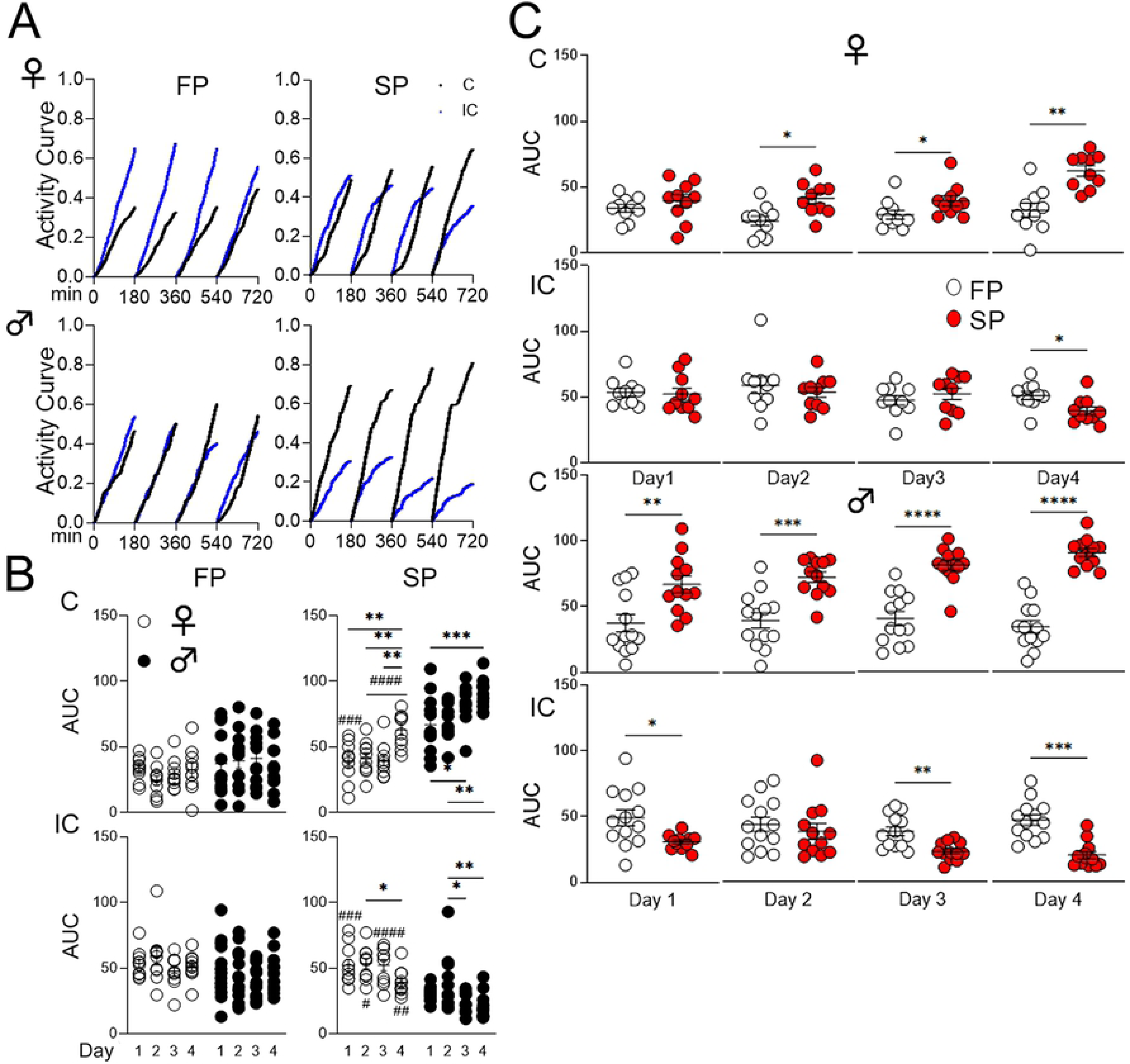
Place learning with corner switch, cohort B. **(A)** Activity curves showing the fractional accumulation (y-axis) of visits over drinking session time (x-axis), reset every 180 minutes, by sex and pass. Curves for correct and incorrect visits are black and blue, respectively. **(B)** Sex comparison of area under the curve (AUC) for activity curves of individual animals by visit category and pass. “*” denotes significant comparisons within sex across program days, while “#” denotes those for the corresponding program day between sexes. Female (♀) and male (♂) data points are indicated by white and black circles, respectively. **(C)** Pass comparison of AUCs for activity curves of individual animals by sex, program day, and visit category. Data points from the first and second passes are indicated by white and red circles, respectively. All data represented as mean ± SEM (\**p* < 0.05, \*\**p* < 0.01, \*\*\**p* < 0.001, \*\*\*\**p* < 0.0001, similarly for ^#^*p* < 0.05, etc.). See Tables 9-11 for statistical analysis. FP = first pass (6-8 weeks), SP = second pass (4-5 months), C = correct (visits), IC = incorrect (visits).

**Table 9.** Statistical analysis of data shown in Figure 6B for place learning with corner switch, cohort B, first pass (6-8 weeks). A *p*-value less than 0.05 is considered significant. ns = not significant.

| Figure 6B | Two-way RM ANOVA, place learning with corner switch, cohort B, first pass |  |  |
| --- | --- | --- | --- |
| Correct<br>(C-AUC) | Source of Variation | F (DFn, DFd) | P |
|  | Interaction | F (3, 63) = 1.196 | 0.3186 |
|  | Day | F (3, 63) = 0.3236 | 0.8083 |
|  | Sex | F (1, 21) = 2.476 | 0.1306 |
| Incorrect<br>(IC-AUC) | Source of Variation | F (DFn, DFd) | P |
|  | Interaction | F (3, 63) = 0.6067 | 0.6131 |
|  | Day | F (3, 63) = 1.309 | 0.2795 |
|  | Sex | F (1, 21) = 4.759 | 0.0406 |
|  | post-hoc Sidak's multiple comp. test | Summary | Adjusted P |
|  | Female vs. Male, day 1 | ns | 0.9552 |
|  | Female vs. Male, day 2 | ns | 0.1174 |
|  | Female vs. Male, day 3 | ns | 0.6359 |
|  | Female vs. Male, day 4 | ns | 0.9713 |

**Table 10.** Statistical analysis of data shown in Figure 6B for place learning with corner switch, cohort B, second pass (4-5 months). A *p*-value less than 0.05 is considered significant (*^/#^*p* < 0.05, **^/##^*p* < 0.01, ***^/###^*p* < 0.001, ^####^*p* < 0.0001). Some non-significant comparisons are omitted. ns = not significant.

| Figure 6B | Two-way RM ANOVA, place learning with corner switch, cohort B, second pass |  |  |
| --- | --- | --- | --- |
| Correct (C-AUC) | Source of Variation | F (DFn, DFd) | P |
|  | Interaction | F (3, 60) = 1.594 | 0.2003 |
|  | Day | F (3, 60) = 13.62 | <0.0001 |
|  | Sex | F (1, 20) = 68.24 | <0.0001 |
|  | post-hoc Sidak's multiple comp. test | Summary | Adjusted P |
|  | Female, day 1 vs. day 4 | ** | 0.0011 |
|  | Female, day 2 vs. day 4 | ** | 0.0039 |
|  | Female, day 3 vs. day 4 | ** | 0.0013 |
|  | Male, day 1 vs. day 2 | ns | 0.8863 |
|  | Male, day 1 vs. day 3 | * | 0.0324 |
|  | Male, day 1 vs. day 4 | *** | 0.0002 |
|  | Male, day 2 vs. day 3 | ns | 0.3527 |
|  | Male, day 2 vs. day 4 | ** | 0.0068 |
|  | Male, day 3 vs. day 4 | ns | 0.5392 |
|  | Female vs. Male, day 1 | ### | 0.0001 |
|  | Female vs. Male, day 2 | #### | <0.0001 |
|  | Female vs. Male, day 3 | #### | <0.0001 |
|  | Female vs. Male, day 4 | #### | <0.0001 |
| Incorrect (IC-AUC) | Source of Variation | F (DFn, DFd) | P |
|  | Interaction | F (3, 60) = 1.427 | 0.2438 |
|  | Day | F (3, 60) = 7.773 | 0.0002 |
|  | Sex | F (1, 20) = 39.84 | <0.0001 |
|  | post-hoc Sidak's multiple comp. test | Summary | Adjusted P |
|  | Female, day 1 vs. day 4 | ns | 0.0821 |
|  | Female, day 2 vs. day 4 | * | 0.0423 |
|  | Female, day 3 vs. day 4 | ns | 0.0773 |
|  | Male, day 1 vs. day 2 | ns | 0.4405 |
|  | Male, day 1 vs. day 3 | ns | 0.5832 |
|  | Male, day 1 vs. day 4 | ns | 0.1874 |
|  | Male, day 2 vs. day 3 | * | 0.0122 |
|  | Male, day 2 vs. day 4 | ** | 0.0016 |
|  | Female vs. Male, day 1 | ### | 0.0006 |
|  | Female vs. Male, day 2 | # | 0.0279 |
|  | Female vs. Male, day 3 | #### | <0.0001 |
|  | Female vs. Male, day 4 | ## | 0.0032 |

**Table 11.** Statistical analysis of data shown in Figure 6C for place learning with corner switch, cohort B, pass comparison. A *p*-value less than 0.05 is considered significant (\**p* < 0.05, \*\**p* < 0.01, \*\*\**p* < 0.001, \*\*\*\**p* < 0.0001). ns = not significant.

| Figure 6C | Paired *t* tests, place learning with corner switch, cohort B, pass comparison |  |  |  |
| --- | --- | --- | --- | --- |
|  | Comparison | *t*, *df* | Summary | *P* |
| Correct<br>(C-AUC) | Female, day 1 | *t*=1.200, *df*=9 | ns | 0.2606 |
|  | Male, day 1 | *t*=3.645, *df*=11 | ** | 0.0039 |
|  | Female, day 2 | *t*=2.577, *df*=9 | * | 0.0298 |
|  | Male, day 2 | *t*=5.053, *df*=11 | *** | 0.0004 |
|  | Female, day 3 | *t*=3.192, *df*=9 | * | 0.0110 |
|  | Male, day 3 | *t*=6.174, *df*=11 | **** | <0.0001 |
|  | Female, day 4 | *t*=4.269, *df*=9 | ** | 0.0021 |
|  | Male, day 4 | *t*=9.086, *df*=11 | **** | <0.0001 |
| Incorrect<br>(IC-AUC) | Comparison | *t*, *df* | Summary | *P* |
|  | Female, day 1 | *t*=0.2539, *df*=9 | ns | 0.8053 |
|  | Male, day 1 | *t*=2.684, *df*=11 | * | 0.0213 |
|  | Female, day 2 | *t*=0.7319, *df*=9 | ns | 0.4828 |
|  | Male, day 2 | *t*=0.8458, *df*=11 | ns | 0.4157 |
|  | Female, day 3 | *t*=0.6925, *df*=9 | ns | 0.5061 |
|  | Male, day 3 | *t*=3.845, *df*=11 | ** | 0.0027 |
|  | Female, day 4 | *t*=2.893, *df*=9 | * | 0.0178 |
|  | Male, day 4 | *t*=4.633, *df*=11 | *** | 0.0007 |

### App^h/h^ rats in cohorts A and B can acquire a behavioral sequencing task by 6-8 weeks of age with session-wise improvement

After place learning and place reversal (cohort A) or place learning with corner switch (cohort B), we further tested the rats’ spatial learning capabilities with a behavioral sequencing program requiring the animals to shuttle between diagonally opposing corners for water access. Visits were categorized as correct (C), lateral (L), or opposite (O) as described in the methods, with activity curves generated (Figures 7A and 8A) and AUC analysis performed (Figures 7B-C and 8B-C) similarly as in other programs. Cohort A rats of both sexes during the first pass showed significant increases in C-AUC and decreases in O-AUC, but no significant changes in L-AUC (Figure 7B). These changes were consistent during the second pass for females, whereas males no longer showed significant session- wise changes in C-AUC (Figure 7B). There were significant sex differences observed for the 3^rd^ drinking session of the first pass for C-AUC (higher in males) and L-AUC (higher in females), with no significant differences observed during the second pass (Figure 7B). For females during the second pass compared to the first, C-AUC was significantly greater for the 1^st^ drinking session; for males, C-AUC was lower while L-AUC was higher for the 3^rd^ drinking session (Figure 7C). Cohort B rats exhibited a different activity profile: for females, the session-wise differences in C-AUC, L-AUC, and O-AUC were minimal during both passes, whereas for males, there were many significant session-wise differences, especially during the second pass with increases in C-AUC and decreases in L-AUC and O-AUC (Figure 8B). Significant differences found between passes for females were sporadic for C-AUC (1^st^ drinking session), L-AUC (4^th^ drinking session), and O-AUC (1^st^ and 3^rd^ drinking sessions); in contrast, for males, C-AUC was significantly higher for every drinking session during the second pass compared to the first, with L-AUC lower for every drinking session except the 1^st^ and O-AUC higher in the 1^st^ and 2^nd^ drinking sessions (Figure 8C).

**Figure 7.**
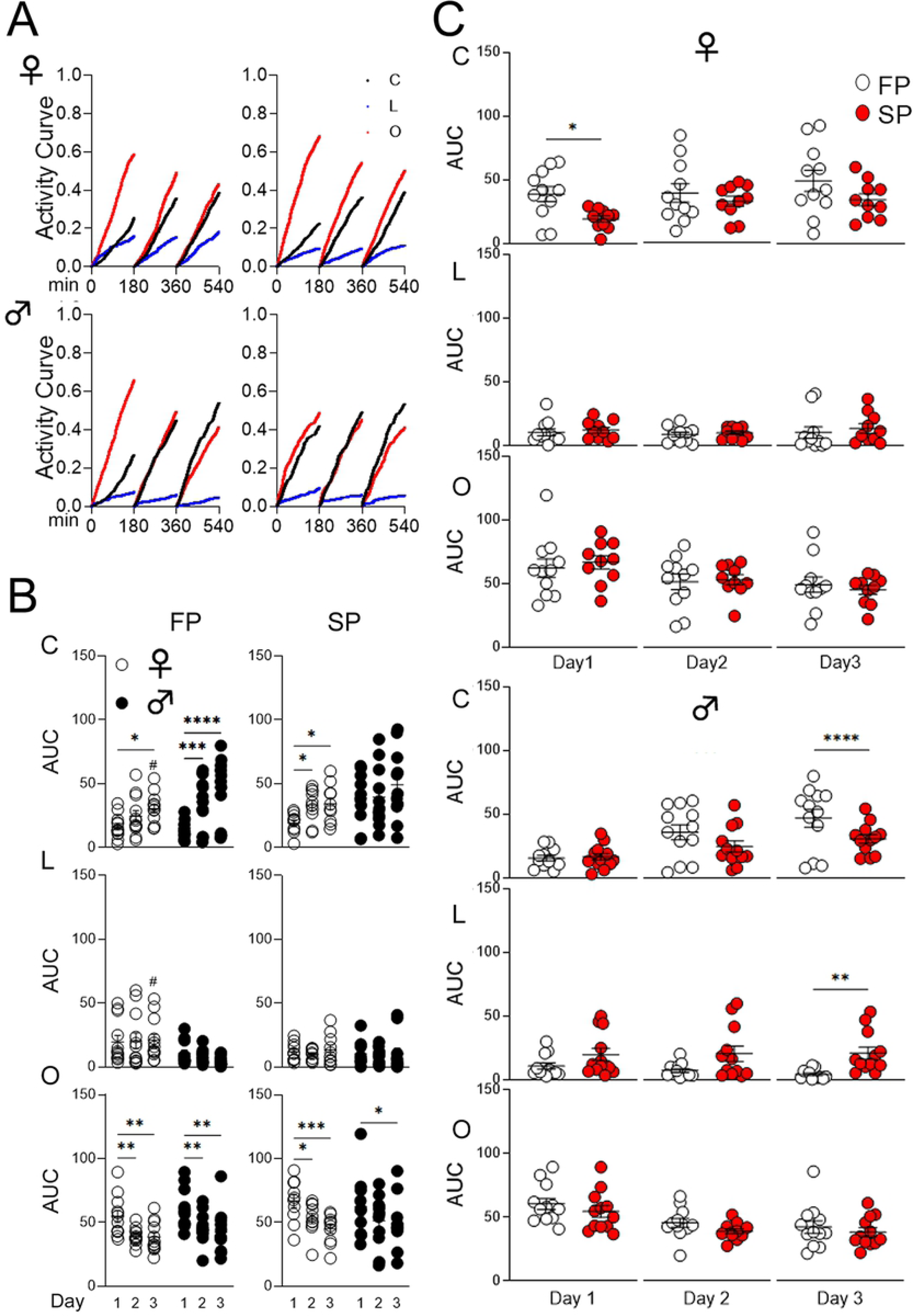
Behavioral sequencing, cohort A. **(A)** Activity curves showing the fractional accumulation (y-axis) of visits over drinking session time (x-axis), reset every 180 minutes, by sex and pass. Curves for correct, lateral, and opposite visits are black, blue, and red, respectively. **(B)** Sex comparison of area under the curve (AUC) for activity curves of individual animals by visit category and pass. “*” denotes significant comparisons within sex across program days, while “#” denotes those for the corresponding program day between sexes. Female (♀) and male (♂) data points are indicated by white and black circles, respectively. **(C)** Pass comparison of AUCs for activity curves of individual animals by sex, program day, and visit category. Data points from the first and second passes are indicated by white and red circles, respectively. All data are represented as mean ± SEM (\**p* < 0.05, \*\**p* < 0.01, \*\*\**p* < 0.001, \*\*\*\**p* < 0.0001, similarly for ^#^*p* < 0.05, etc.). See Tables 12-14 for statistical analysis. FP = first pass (6-8 weeks), SP = second pass (4-5 months), C = correct (visits), L = lateral (visits), O = opposite (visits).

**Figure 8.**
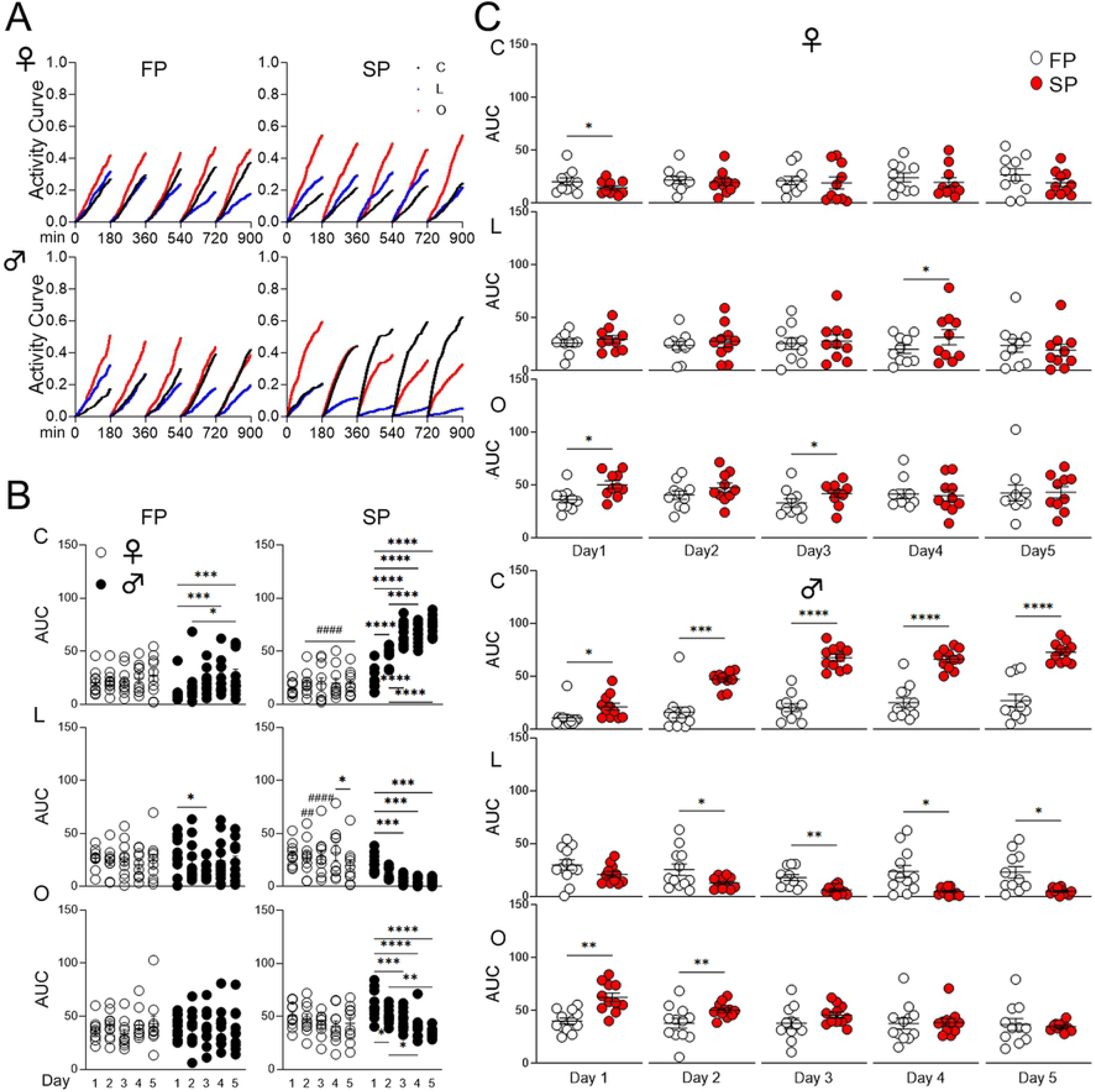
Behavioral sequencing, cohort B. **(A)** Activity curves showing the fractional accumulation (y-axis) of visits over drinking session time (x-axis), reset every 180 minutes, by sex and pass. Curves for correct, lateral, and opposite visits are black, blue, and red, respectively. **(B)** Sex comparison of area under the curve (AUC) for activity curves of individual animals by visit category and pass. “*” denotes significant comparisons within sex across program days, while “#” denotes those for the corresponding program day between sexes. Female (♀) and male (♂) data points are indicated by white and black circles, respectively. **(C)** Pass comparison of AUCs for activity curves of individual animals by sex, program day, and visit category. Data points from the first and second passes are indicated by white and red circles, respectively. All data are represented as mean ± SEM (\**p* < 0.05, \*\**p* < 0.01, \*\*\**p* < 0.001, \*\*\*\**p* < 0.0001, similarly for ^#^*p* < 0.05, etc.). See Tables 15-17 for statistical analysis. FP = first pass (6-8 weeks), SP = second pass (4-5 months), C = correct (visits), L = lateral (visits), O = opposite (visits).

**Table 12.** – Statistical analysis of data shown in Figure 7B for behavioral sequencing, cohort A, first pass (6-8 weeks). A *p*-value less than 0.05 is considered significant (*^/#^*p* < 0.05, \*\**p* < 0.01, \*\*\**p* < 0.001, \*\*\*\**p* < 0.0001). ns = not significant.

| **Figure 7B** |  |  |  |
| --- | --- | --- | --- |
| **Two-way RM ANOVA, behavioral sequencing, cohort A, first pass** |  |  |  |
| Correct<br>(C-AUC) | **Source of Variation** | **F (DFn, DFd)** | **P** |
|  | Interaction | F (2, 44) = 3.323 | 0.0453 |
|  | Day | F (2, 44) = 22.41 | <0.0001 |
|  | Sex | F (1, 22) = 2.906 | 0.1023 |
|  | **post-hoc Sidak's multiple comp. test** | **Summary** | **Adjusted P** |
|  | Female, day 1 vs. day 2 | ns | 0.2721 |
|  | Female, day 1 vs. day 3 | * | 0.0168 |
|  | Female, day 2 vs. day 3 | ns | 0.5330 |
|  | Male, day 1 vs. day 2 | *** | 0.0004 |
|  | Male, day 1 vs. day 3 | **** | <0.0001 |
|  | Male, day 2 vs. day 3 | ns | 0.0775 |
| Lateral<br>(L-AUC) | **Source of Variation** | **F (DFn, DFd)** | **P** |
|  | Interaction | F (2, 44) = 1.391 | 0.2595 |
|  | Day | F (2, 44) = 0.6175 | 0.5439 |
|  | Sex | F (1, 22) = 7.170 | 0.0137 |
|  | **post-hoc Sidak's multiple comp. test** | **Summary** | **Adjusted P** |
|  | Female vs. Male, day 1 | ns | 0.2961 |
|  | Female vs. Male, day 2 | ns | 0.0578 |
|  | Female vs. Male, day 3 | # | 0.0109 |
| Opposite<br>(O-AUC) | **Source of Variation** | **F (DFn, DFd)** | **P** |
|  | Interaction | F (2, 44) = 0.09638 | 0.9083 |
|  | Day | F (2, 44) = 15.98 | <0.0001 |
|  | Sex | F (1, 22) = 2.116 | 0.1599 |
|  | **post-hoc Sidak's multiple comp. test** | **Summary** | **Adjusted P** |
|  | Female, day 1 vs. day 2 | ** | 0.0055 |
|  | Female, day 1 vs. day 3 | ** | 0.0039 |
|  | Female, day 2 vs. day 3 | ns | 0.9992 |
|  | Male, day 1 vs. day 2 | ** | 0.0088 |
|  | Male, day 1 vs. day 3 | ** | 0.0011 |
|  | Male, day 2 vs. day 3 | ns | 0.8555 |

**Table 13.**
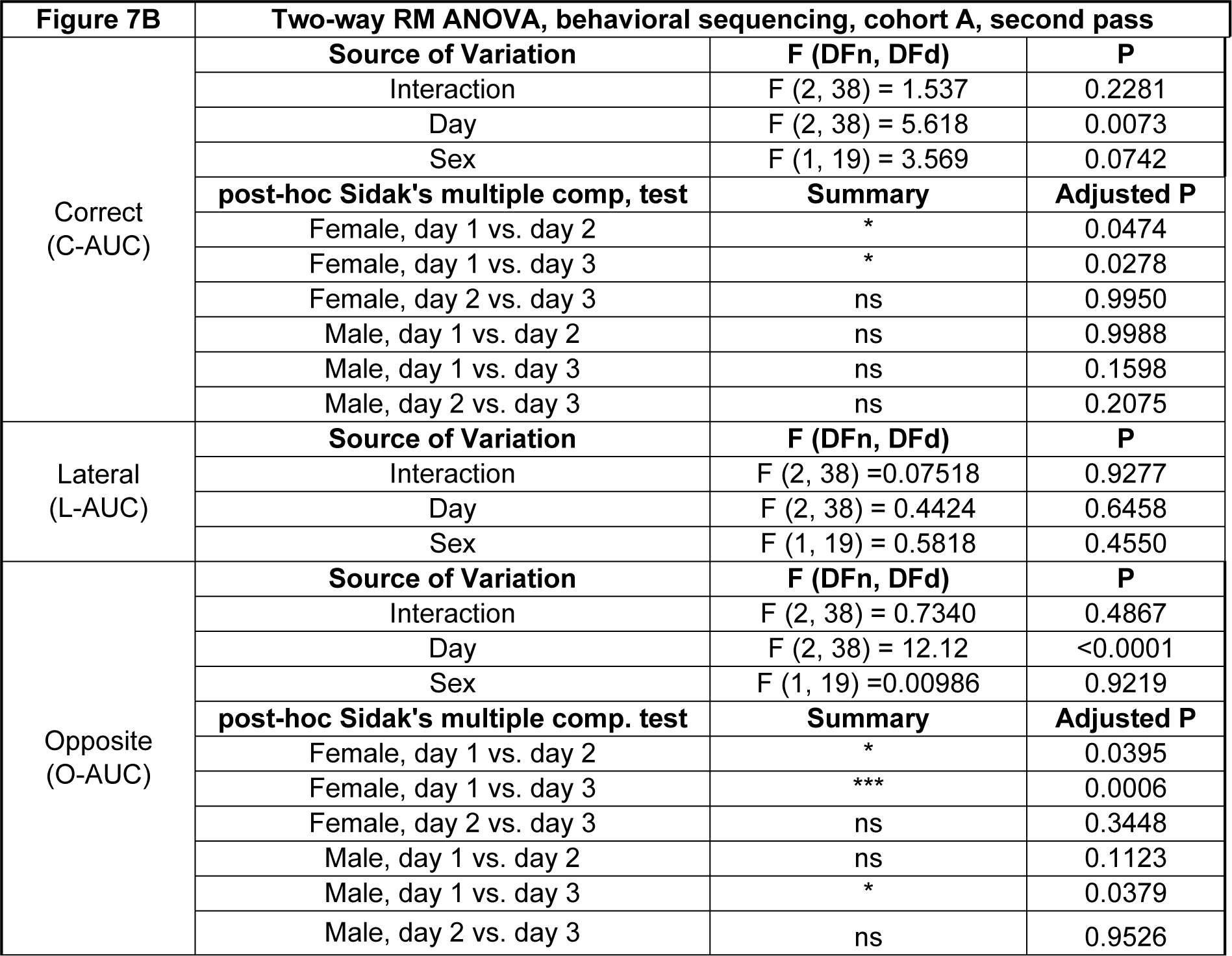
Statistical analysis of data shown in Figure 7B for behavioral sequencing, cohort A, second pass (4-5 months). A *p*-value less than 0.05 is considered significant (\**p* < 0.05, \*\*\**p* < 0.001). ns = not significant.

**Table 14.** – Statistical analysis of data shown in Figure 7C for behavioral sequencing, cohort A, pass comparison. A *p*-value less than 0.05 is considered significant (\**p* < 0.05, \*\**p* < 0.01). ns = not significant.

| **Figure 7C** | **Paired *t* tests, behavioral sequencing, cohort A, pass comparison** |  |  |  |
| --- | --- | --- | --- | --- |
|  | **Comparison** | **t, df** | **Summary** | **P** |
| Correct<br>(C-AUC) | Female, day 1 | t=2.491, df=8 | * | 0.0375 |
|  | Male, day 1 | t=0.2282, df=11 | ns | 0.8237 |
|  | Female, day 2 | t=1.032, df=8 | ns | 0.3324 |
|  | Male, day 2 | t=1.474, df=11 | ns | 0.1686 |
|  | Female, day 3 | t=1.177, df=8 | ns | 0.2728 |
|  | Male, day 3 | t=1.844, df=11 | ns | 0.0923 |
| Lateral<br>(L-AUC) | **Comparison** | **t, df** | **Summary** | **P** |
|  | Female, day 1 | t=0.9930, df=8 | ns | 0.3498 |
|  | Male, day 1 | t=1.502, df=11 | ns | 0.1613 |
|  | Female, day 2 | t=1.498, df=8 | ns | 0.1724 |
|  | Male, day 2 | t=2.034, df=11 | ns | 0.0668 |
|  | Female, day 3 | t=0.3042, df=8 | ns | 0.7687 |
|  | Male, day 3 | t=3.328, df=11 | ** | 0.0067 |
| Opposite<br>(O-AUC) | **Comparison** | **t, df** | **Summary** | **P** |
|  | Female, day 1 | t=0.9581, df=8 | ns | 0.3661 |
|  | Male, day 1 | t=0.9490, df=11 | ns | 0.3630 |
|  | Female, day 2 | t=0.6758, df=8 | ns | 0.5182 |
|  | Male, day 2 | t=1.496, df=11 | ns | 0.1628 |
|  | Female, day 3 | t=0.4350, df=8 | ns | 0.6751 |
|  | Male, day 3 | t=0.7209, df=11 | ns | 0.4860 |

**Table 15.** Statistical analysis of data shown in Figure 8B for behavioral sequencing, cohort B, first pass (6-8 weeks). A *p*-value less than 0.05 is considered significant (\**p* < 0.05, \*\*\**p* < 0.001). ns = not significant.

| Figure 8B | Mixed-effects analysis, behavioral sequencing, cohort B, first pass |  |  |
| --- | --- | --- | --- |
| Correct<br>(C-AUC) | Source of Variation | F (DFn, DFd) | P |
|  | Interaction | F (4, 77) = 1.576 | 0.1891 |
|  | Day | F (4, 77) = 6.336 | 0.0002 |
|  | Sex | F (1, 21) = 0.5258 | 0.4764 |
|  | post-hoc Sidak's multiple comp. test | Summary | Adjusted P |
|  | Female, day 1 vs. day 2 | ns | >0.9999 |
|  | Female, day 1 vs. day 3 | ns | >0.9999 |
|  | Female, day 1 vs. day 4 | ns | 0.9681 |
|  | Female, day 1 vs. day 5 | ns | 0.5746 |
|  | Female, day 2 vs. day 3 | ns | >0.9999 |
|  | Female, day 2 vs. day 4 | ns | 0.9994 |
|  | Female, day 2 vs. day 5 | ns | 0.8721 |
|  | Female, day 3 vs. day 4 | ns | 0.9980 |
|  | Female, day 3 vs. day 5 | ns | 0.8138 |
|  | Female, day 4 vs. day 5 | ns | 0.9987 |
|  | Male, day 1 vs. day 2 | ns | 0.6038 |
|  | Male, day 1 vs. day 3 | ns | 0.0745 |
|  | Male, day 1 vs. day 4 | *** | 0.0004 |
|  | Male, day 1 vs. day 5 | *** | 0.0002 |
|  | Male, day 2 vs. day 3 | ns | 0.9644 |
|  | Male, day 2 vs. day 4 | ns | 0.0898 |
|  | Male, day 2 vs. day 5 | * | 0.0388 |
|  | Male, day 3 vs. day 4 | ns | 0.7651 |
|  | Male, day 3 vs. day 5 | ns | 0.4960 |
|  | Male, day 4 vs. day 5 | ns | >0.9999 |
| Lateral<br>(L-AUC) | Source of Variation | F (DFn, DFd) | P |
|  | Interaction | F (4, 77) = 1.215 | 0.3114 |
|  | Day | F (4, 77) = 1.408 | 0.2394 |
|  | Sex | F (1, 21) = 0.0001 | 0.9910 |
| Opposite<br>(O-AUC) | Source of Variation | F (DFn, DFd) | P |
|  | Interaction | F (4, 77) = 0.7529 | 0.5592 |
|  | Day | F (4, 77) = 0.4977 | 0.7374 |
|  | Sex | F (1, 21) = 0.0369 | 0.8496 |

**Table 16.** Statistical analysis of data shown in Figure 8B for behavioral sequencing, cohort B, second pass (4-5 months). A *p*-value less than 0.05 is considered significant (\**p* < 0.05, ^##^*p* < 0.01, \*\*\**p* < 0.001, ****^/####^*p* < 0.0001). Non-significant comparisons have been omitted.

| Figure 8B | Two-way RM ANOVA, behavioral sequencing, cohort B, second pass |  |  |
| --- | --- | --- | --- |
| Correct<br>(C-AUC) | Source of Variation | F (DFn, DFd) | P |
|  | Interaction | F (4, 76) = 32.02 | <0.0001 |
|  | Day | F (4, 76) = 46.43 | <0.0001 |
|  | Sex | F (1, 19) = 84.69 | <0.0001 |
|  | post-hoc Sidak's multiple comparisons test | Summary | Adjusted P |
|  | Male, day 1 vs. days 2, 3, 4, 5 | **** | <0.0001 |
|  | Male, day 2 vs. day 3 | **** | <0.0001 |
|  | Male, day 2 vs. day 4 | **** | <0.0001 |
|  | Male, day 2 vs. day 5 | **** | <0.0001 |
|  | Female vs Male, day 2 | #### | <0.0001 |
|  | Female vs Male, day 3 | #### | <0.0001 |
|  | Female vs Male, day 4 | #### | <0.0001 |
|  | Female vs Male, day 5 | #### | <0.0001 |
| Lateral<br>(L-AUC) | Source of Variation | F (DFn, DFd) | P |
|  | Interaction | F (4, 76) = 3.906 | 0.0061 |
|  | Day | F (4, 76) = 7.183 | <0.0001 |
|  | Sex | F (1, 19) = 13.76 | 0.0015 |
|  | post-hoc Sidak's multiple comparisons test | Summary | Adjusted P |
|  | Female, day 4 vs. day 5 | * | 0.0155 |
|  | Male, day 1 vs. day 3 | *** | 0.0006 |
|  | Male, day 1 vs. day 4 | *** | 0.0001 |
|  | Male, day 1 vs. day 5 | *** | 0.0001 |
|  | Female vs. male, day 3 | ## | 0.0011 |
|  | Female vs. male, day 4 | #### | <0.0001 |
| Opposite<br>(O-AUC) | Source of Variation | F (DFn, DFd) | P |
|  | Interaction | F (4, 76) = 3.607 | 0.0096 |
|  | Day | F (4, 76) = 13.74 | <0.0001 |
|  | Sex | F (1, 19) = 0.1984 | 0.6611 |
|  | post-hoc Sidak's multiple comparisons test | Summary | Adjusted P |
|  | Male, day 1 vs. day 2 | * | 0.0227 |
|  | Male, day 1 vs. day 3 | *** | 0.0004 |
|  | Male, day 1 vs. day 4 | **** | <0.0001 |
|  | Male, day 1 vs. day 5 | **** | <0.0001 |
|  | Male, day 2 vs. day 4 | * | 0.0307 |
|  | Male, day 2 vs. day 5 | ** | 0.0014 |

**Table 17.**
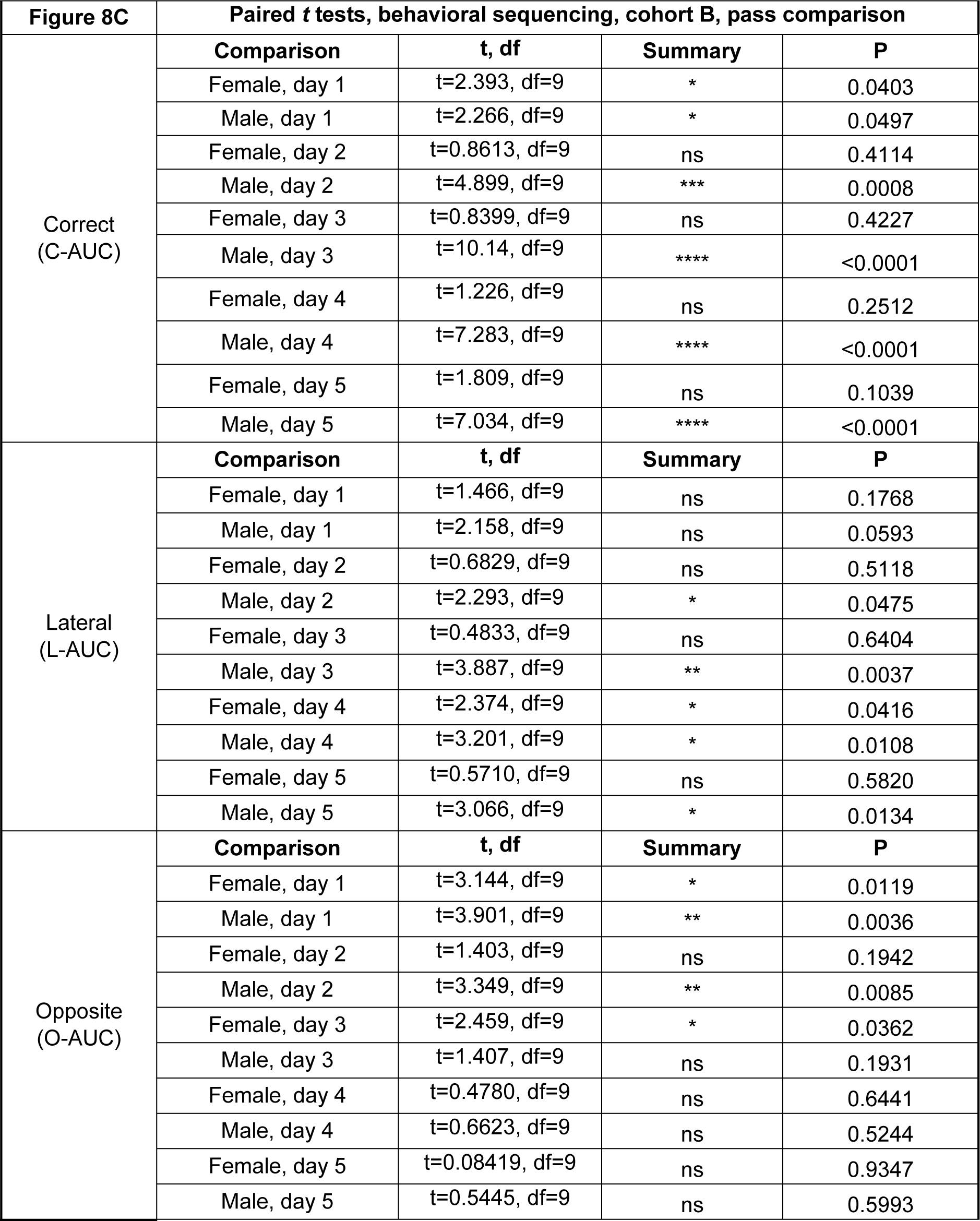
Statistical analysis of data shown in Figure 8C for behavioral sequencing, cohort B, pass comparison. A *p*-value less than 0.05 is considered significant (\**p* < 0.05, \*\**p* < 0.01, \*\*\**p* < 0.001, \*\*\*\**p* < 0.0001). ns = not significant.

### *App*^h/h^ rats in cohort A and B may not be able to acquire a serial reversal task by 4-5 months of age

We ended the timeline for both cohorts with a serial reversal program designed to add a layer of complexity to behavioral sequencing by requiring the rats to alternate diagonals after every eight correct nosepokes. For cohort A, qualitatively, activity curves for both sexes did not show much difference between passes or session-wise improvement (Figure 9A). Session-wise differences in AUC were minimal for both sexes during both passes, with some significant sex differences during the second pass (Figure 9B). Significant differences between passes were sporadic for females in L-AUC (2^nd^ drinking session) and O-AUC (1^st^ drinking session), and non-existent for males (Figure 9C). For cohort B, the activity curves show a possible difference between the first and second pass for males, but no session-wise differences (Figure 10A). AUC analysis revealed that for males compared to females during the second pass, C-AUC was significantly higher for all drinking sessions, with O-AUC significantly lower for the 3^rd^ and 4^th^ drinking sessions (Figure 10B). For males during the second pass compared to the first, C-AUC was significantly higher for every drinking session, with L-AUC significantly higher for the 4^th^ drinking session; significant differences between passes were non- existent for females (Figure 10C).

**Figure 9.**
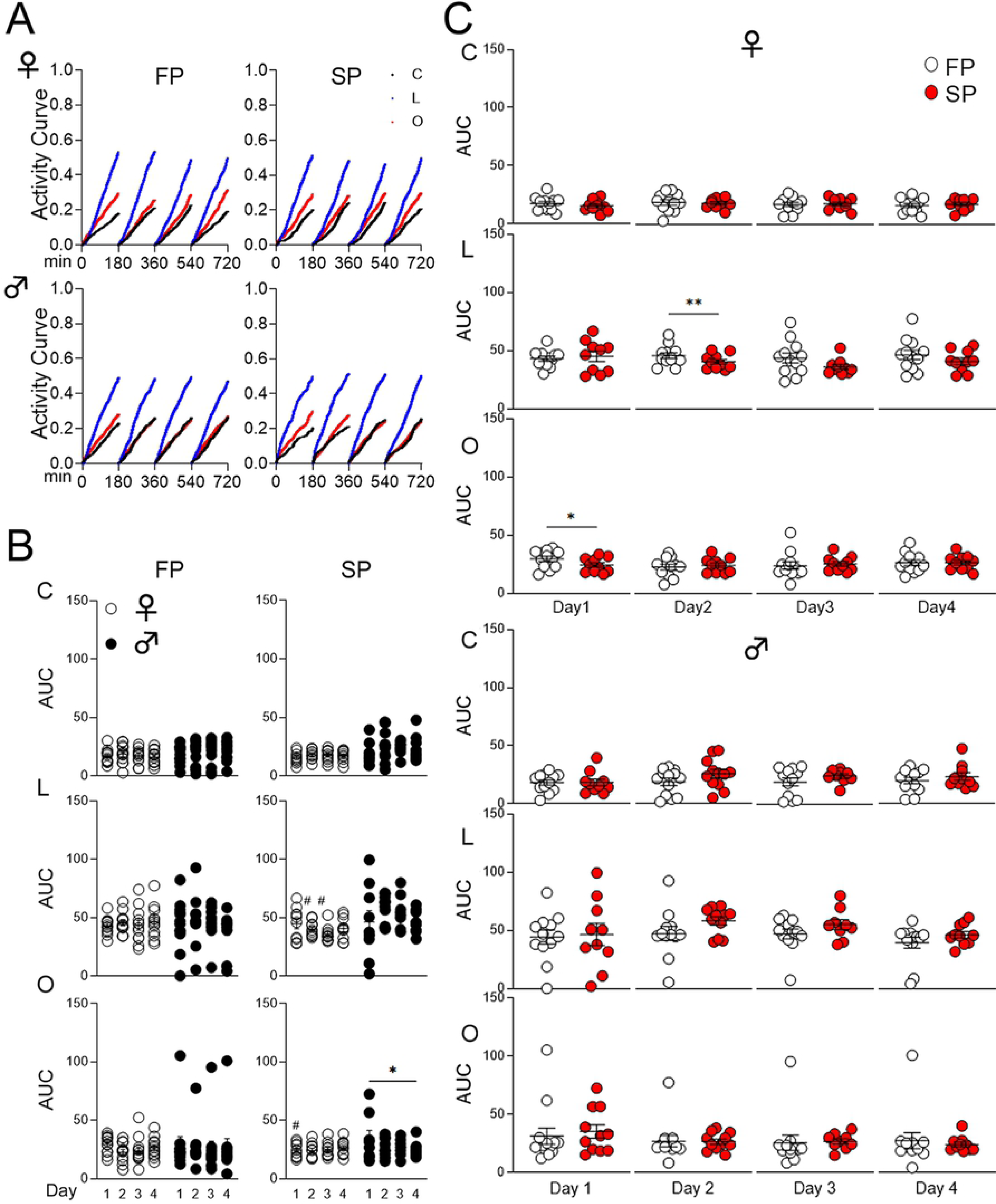
Serial reversal, cohort A. **(A)** Activity curves showing the fractional accumulation (y-axis) of visits over drinking session time (x-axis), reset every 180 minutes, by sex and pass. Curves for correct, lateral, and opposite visits are black, blue, and red, respectively. **(B)** Sex comparison of area under the curve (AUC) for activity curves of individual animals by visit category and pass. “*” denotes significant comparisons within sex across program days, while “#” denotes those for the corresponding program day between sexes. Female (♀) and male (♂) data points are indicated by white and black circles, respectively. **(C)** Pass comparison of AUCs for activity curves of individual animals by sex, program day, and visit category. Data points from the first and second passes are indicated by white and red circles, respectively. All data are represented as mean ± SEM (\**p* < 0.05, \*\**p* < 0.01, \*\*\**p* < 0.001, \*\*\*\**p* < 0.0001, similarly for ^#^*p* < 0.05, etc.). See Tables 18-20 for statistical analysis. FP = first pass (6-8 weeks), SP = second pass (4-5 months), C = correct (visits), L = lateral (visits), O = opposite (visits).

**Figure 10.**
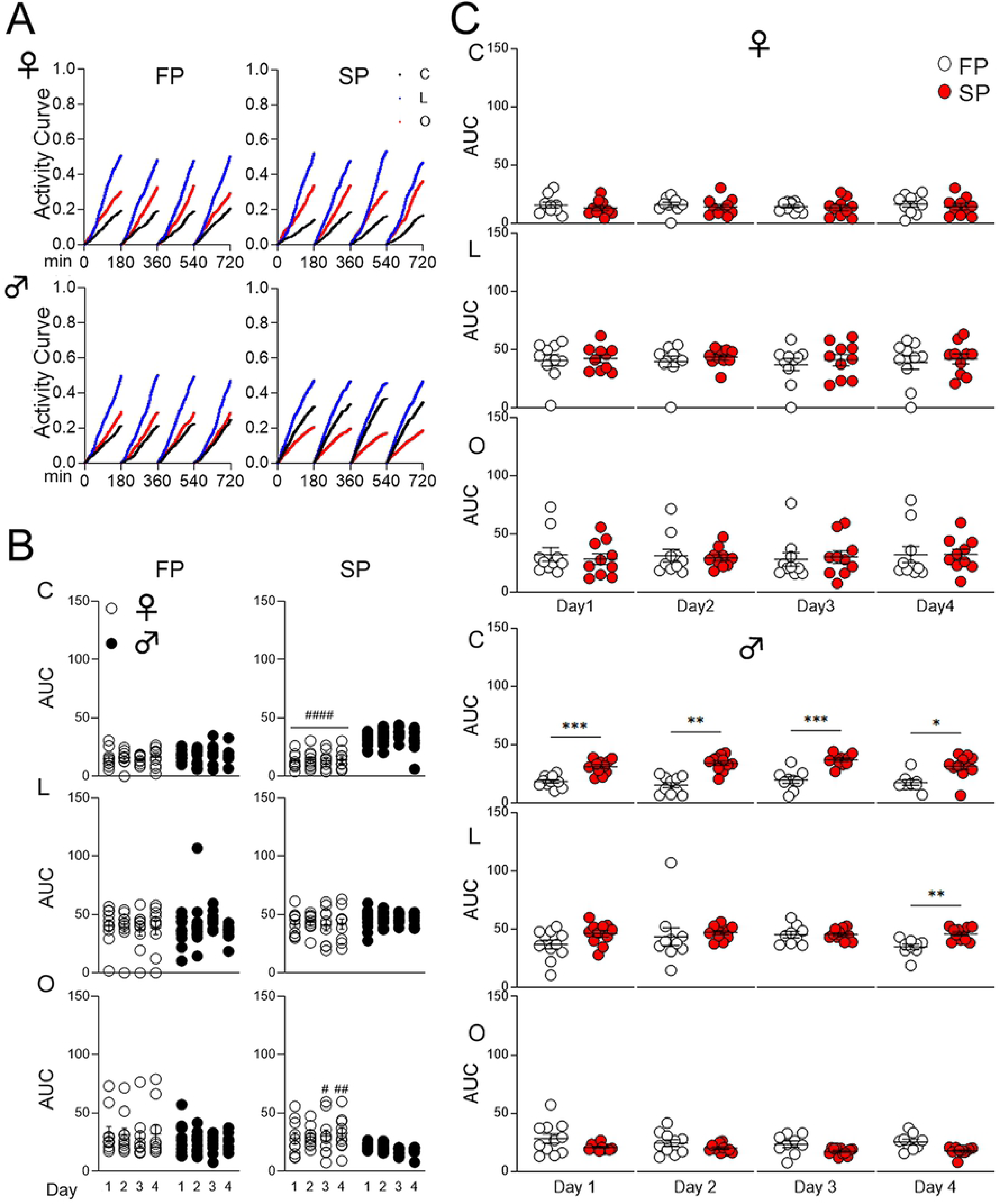
Serial reversal, cohort B. **(A)** Activity curves showing the fractional accumulation (y-axis) of visits over drinking session time (x-axis), reset every 180 minutes, by sex and pass. Curves for correct, lateral, and opposite visits are black, blue, and red, respectively. **(B)** Sex comparison of area under the curve (AUC) for activity curves of individual animals by visit category and pass. “*” denotes significant comparisons within sex across program days, while “#” denotes those for the corresponding program day between sexes. Female (♀) and male (♂) data points are indicated by white and black circles, respectively. **(C)** Pass comparison of AUCs for activity curves of individual animals by sex, program day and visit category. Data points from the first and second passes are indicated by white and red circles, respectively. All data are represented as mean ± SEM (\**p* < 0.05, \*\**p* < 0.01, \*\*\**p* < 0.001, \*\*\*\**p* < 0.0001, similarly for ^#^*p* < 0.05, etc.). See Tables 21-23 for statistical analysis. FP = first pass (6-8 weeks), SP = second pass (4-5 months), C = correct (visits), L = lateral (visits), O = opposite (visits).

**Table 18.**
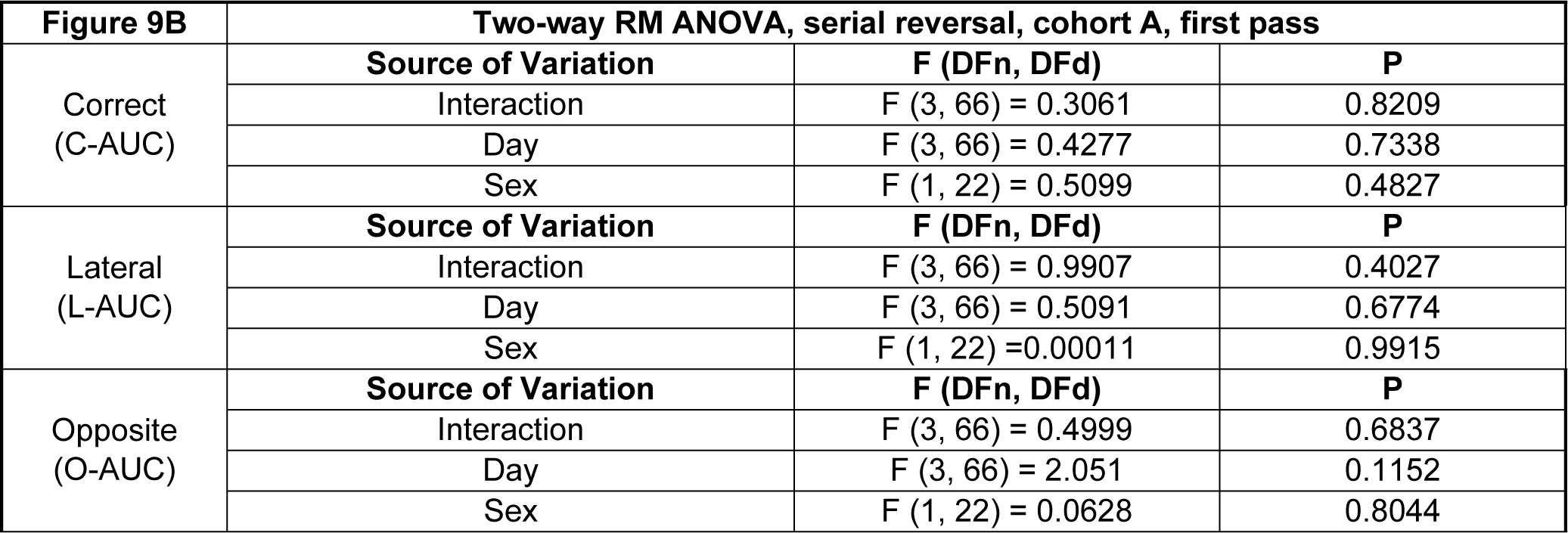
Statistical analysis of data shown in Figure 9B for serial reversal, cohort A, first pass (6-8 weeks). A *p*-value less than 0.05 is considered significant.

| Figure 9B Two-way RM ANOVA, serial reversal, cohort A, first pass |  |  |  |
| --- | --- | --- | --- |
|  | Source of Variation | F (DFn, DFd) | P |
| Correct (C-AUC) | Interaction | F (3, 66) = 0.3061 | 0.8209 |
|  | Day | F (3, 66) = 0.4277 | 0.7338 |
|  | Sex | F (1, 22) = 0.5099 | 0.4827 |
| Lateral (L-AUC) | Source of Variation | F (DFn, DFd) | P |
|  | Interaction | F (3, 66) = 0.9907 | 0.4027 |
|  | Day | F (3, 66) = 0.5091 | 0.6774 |
|  | Sex | F (1, 22) = 0.00011 | 0.9915 |
| Opposite (O-AUC) | Source of Variation | F (DFn, DFd) | P |
|  | Interaction | F (3, 66) = 0.4999 | 0.6837 |
|  | Day | F (3, 66) = 2.051 | 0.1152 |
|  | Sex | F (1, 22) = 0.0628 | 0.8044 |

**Table 19.** Statistical analysis of data shown in Figure 9B for serial reversal, cohort A, second pass (4-5 months). A *p*-value less than 0.05 is considered significant (^#^*p* < 0.05). ns = not significant.

| **Figure 9B** | **Mixed-effects analysis, serial reversal, cohort A, second pass** |  |  |
| --- | --- | --- | --- |
| Correct<br>(C-AUC) | **Source of Variation** | **F (DFn, DFd)** | **P** |
|  | Interaction | F (3, 53) = 0.6830 | 0.5664 |
|  | Day | F (3, 53) = 2.568 | 0.0641 |
|  | Sex | F (1, 19) = 5.754 | 0.0269 |
|  | **post-hoc Sidak's multiple comp. test** | **Summary** | **Adjusted P** |
|  | Female vs. Male, day 1 | ns | 0.8521 |
|  | Female vs. Male, day 2 | ns | 0.1465 |
|  | Female vs. Male, day 3 | ns | 0.1190 |
|  | Female vs. Male, day 4 | ns | 0.1574 |
| Lateral<br>(L-AUC) | **Source of Variation** | **F (DFn, DFd)** | **P** |
|  | Interaction | F (3, 53) = 1.923 | 0.1370 |
|  | Day | F (3, 53) = 0.5768 | 0.6328 |
|  | Sex | F (1, 19) = 10.73 | 0.0040 |
|  | **post-hoc Sidak's multiple comp. test** | **Summary** | **Adjusted P** |
|  | Female vs. male, day 1 | ns | 0.9988 |
|  | Female vs. male, day 2 | # | 0.0228 |
|  | Female vs. male, day 3 | # | 0.0203 |
|  | Female vs. male, day 4 | ns | 0.8612 |
| Opposite<br>(O-AUC) | **Source of Variation** | **F (DFn, DFd)** | **P** |
|  | Interaction | F (3, 53) = 2.117 | 0.1090 |
|  | Day | F (3, 53) = 1.178 | 0.3271 |
|  | Sex | F (1, 19) = 1.600 | 0.2212 |

**Table 20.**
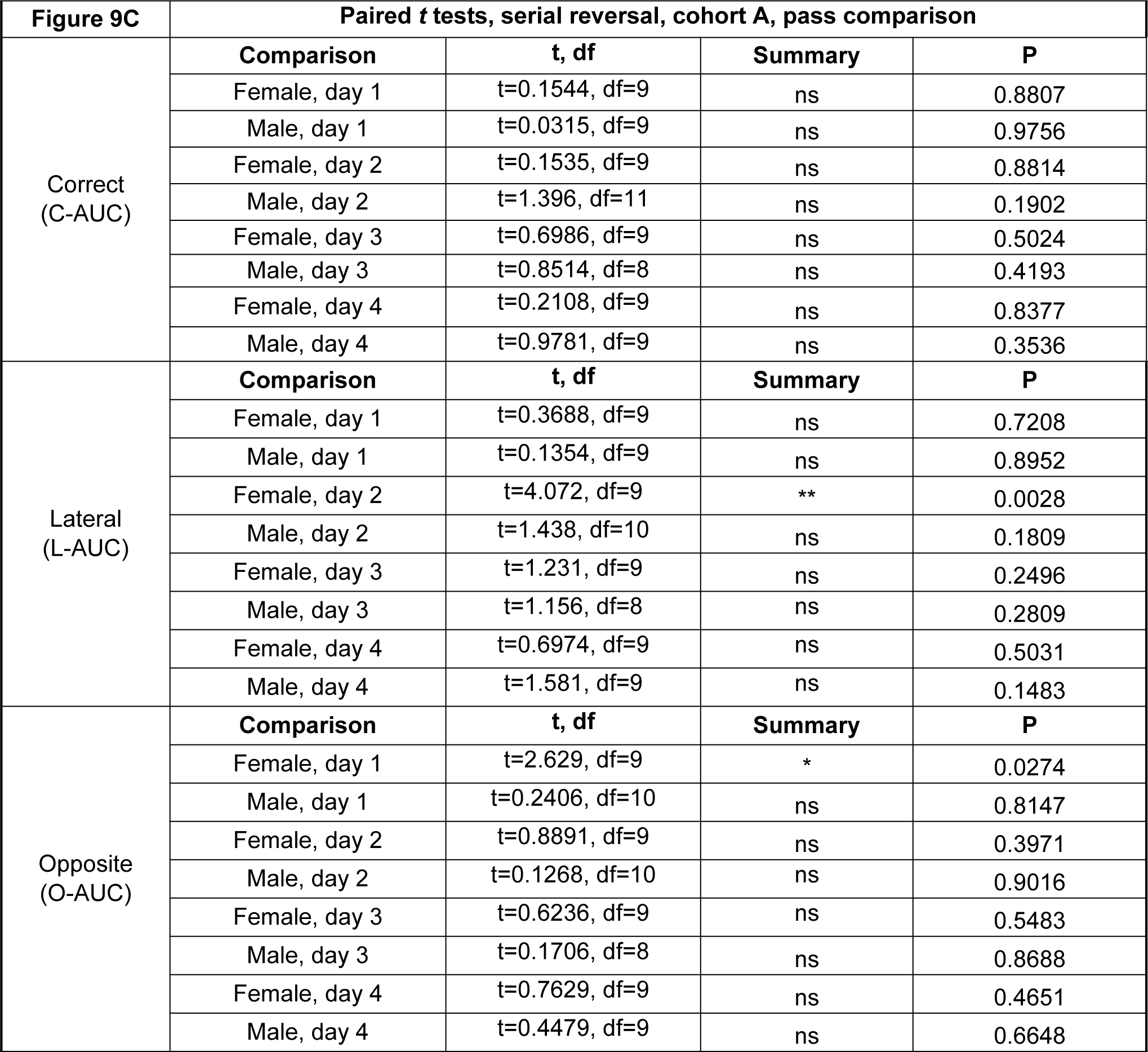
Statistical analysis of data shown in Figure 7C for serial reversal, cohort A, pass comparison. A *p*-value less than 0.05 is considered significant (\**p* < 0.05, \*\**p* < 0.01). ns = not significant.

| **Figure 9C** | **Paired *t* tests, serial reversal, cohort A, pass comparison** |  |  |  |
| --- | --- | --- | --- | --- |
| Correct<br>(C-AUC) | **Comparison** | **t, df** | **Summary** | **P** |
|  | Female, day 1 | t=0.1544, df=9 | ns | 0.8807 |
|  | Male, day 1 | t=0.0315, df=9 | ns | 0.9756 |
|  | Female, day 2 | t=0.1535, df=9 | ns | 0.8814 |
|  | Male, day 2 | t=1.396, df=11 | ns | 0.1902 |
|  | Female, day 3 | t=0.6986, df=9 | ns | 0.5024 |
|  | Male, day 3 | t=0.8514, df=8 | ns | 0.4193 |
|  | Female, day 4 | t=0.2108, df=9 | ns | 0.8377 |
|  | Male, day 4 | t=0.9781, df=9 | ns | 0.3536 |
| Lateral<br>(L-AUC) | **Comparison** | **t, df** | **Summary** | **P** |
|  | Female, day 1 | t=0.3688, df=9 | ns | 0.7208 |
|  | Male, day 1 | t=0.1354, df=9 | ns | 0.8952 |
|  | Female, day 2 | t=4.072, df=9 | ** | 0.0028 |
|  | Male, day 2 | t=1.438, df=10 | ns | 0.1809 |
|  | Female, day 3 | t=1.231, df=9 | ns | 0.2496 |
|  | Male, day 3 | t=1.156, df=8 | ns | 0.2809 |
|  | Female, day 4 | t=0.6974, df=9 | ns | 0.5031 |
|  | Male, day 4 | t=1.581, df=9 | ns | 0.1483 |
| Opposite<br>(O-AUC) | **Comparison** | **t, df** | **Summary** | **P** |
|  | Female, day 1 | t=2.629, df=9 | * | 0.0274 |
|  | Male, day 1 | t=0.2406, df=10 | ns | 0.8147 |
|  | Female, day 2 | t=0.8891, df=9 | ns | 0.3971 |
|  | Male, day 2 | t=0.1268, df=10 | ns | 0.9016 |
|  | Female, day 3 | t=0.6236, df=9 | ns | 0.5483 |
|  | Male, day 3 | t=0.1706, df=8 | ns | 0.8688 |
|  | Female, day 4 | t=0.7629, df=9 | ns | 0.4651 |
|  | Male, day 4 | t=0.4479, df=9 | ns | 0.6648 |

**Table 21.**
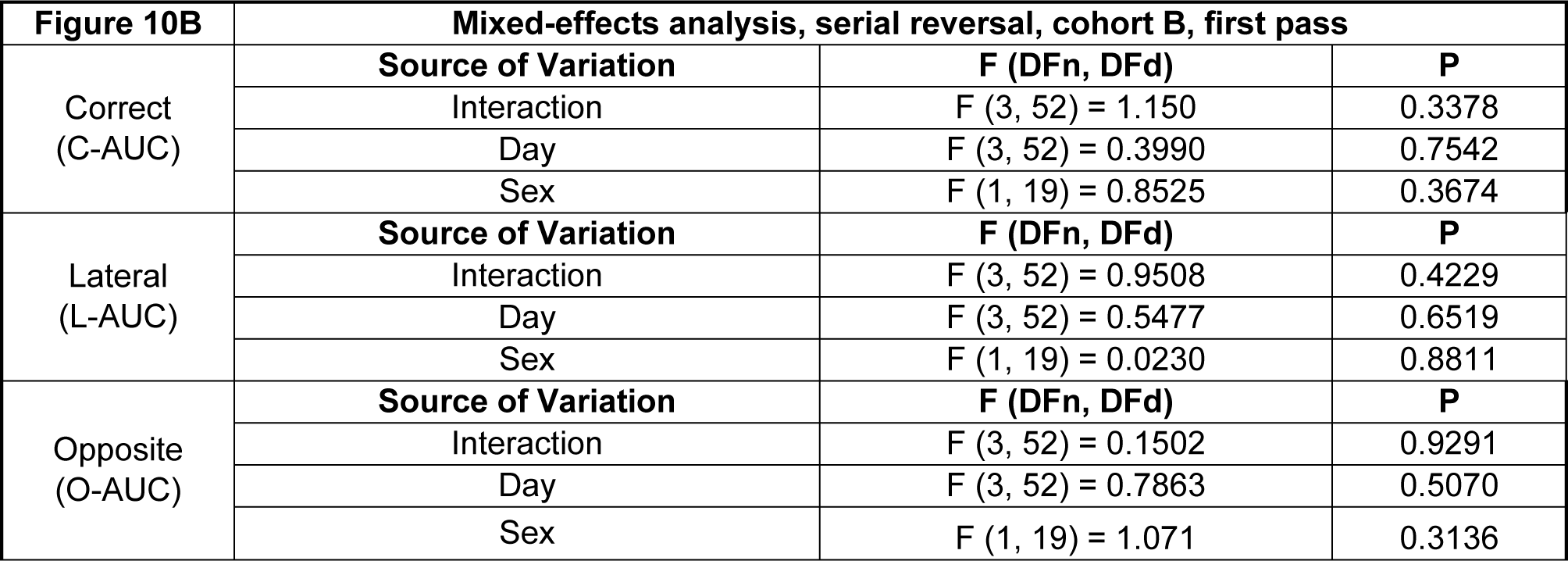
Statistical analysis of data shown in Figure 10B for serial reversal, cohort B, first pass (6-8 weeks). A *p*-value less than 0.05 is considered significant.

**Table 22.** Statistical analysis of data shown in Figure 10B for serial reversal, cohort B, second pass (4-5 months). A *p*-value less than 0.05 is considered significant (^#^*p* < 0.05, ^##^*p* < 0.01, ^####^*p* < 0.0001). ns = not significant.

| **Figure 10B** | **Two-way RM ANOVA, serial reversal, cohort B, second pass** |  |  |
| --- | --- | --- | --- |
|  | **Source of Variation** | **F (DFn, DFd)** | **P** |
| Correct<br>(C-AUC) | Interaction | F (3, 57) = 1.357 | 0.2652 |
|  | Day | F (3, 57) = 1.391 | 0.2548 |
|  | Sex | F (1, 19) = 69.20 | <0.0001 |
|  | **post-hoc Sidak's multiple comp. test** | **Summary** | **Adjusted P** |
|  | Female vs. Male, day 1 | #### | <0.0001 |
|  | Female vs. Male, day 2 | #### | <0.0001 |
|  | Female vs. Male, day 3 | #### | <0.0001 |
|  | Female vs. Male, day 4 | #### | <0.0001 |
| Lateral<br>(L-AUC) | **Source of Variation** | **F (DFn, DFd)** | **P** |
|  | Interaction | F (3, 57) = 0.0096 | 0.9987 |
|  | Day | F (3, 57) = 0.1552 | 0.9259 |
|  | Sex | F (1, 19) = 2.888 | 0.1056 |
| Opposite<br>(O-AUC) | **Source of Variation** | **F (DFn, DFd)** | **P** |
|  | Interaction | F (3, 57) = 0.6388 | 0.5932 |
|  | Day | F (3, 57) = 0.0676 | 0.9769 |
|  | Sex | F (1, 19) = 20.49 | 0.0002 |
|  | **post-hoc Sidak's multiple comp. test** | **Summary** | **Adjusted P** |
|  | Female vs. Male, day 1 | ns | 0.2627 |
|  | Female vs. Male, day 2 | ns | 0.1193 |
|  | Female vs. Male, day 3 | # | 0.0146 |
|  | Female vs. Male, day 4 | ## | 0.0034 |

**Table 23.** Statistical analysis of data shown in Figure 10C for serial reversal, cohort B, pass comparison. A *p*-value less than 0.05 is considered significant (\**p* < 0.05, \*\**p* < 0.01, \*\*\**p* < 0.001). ns = not significant.

| Figure 10C | Paired *t* tests, serial reversal, cohort B, pass comparison |  |  |  |
| --- | --- | --- | --- | --- |
| Correct<br>(C-AUC) | Comparison | *t*, *df* | Summary | *P* |
|  | Female, day 1 | *t*=0.8066, *df*=9 | ns | 0.4407 |
|  | Male, day 1 | *t*=5.117, *df*=9 | *** | 0.0006 |
|  | Female, day 2 | *t*=0.5204, *df*=9 | ns | 0.6153 |
|  | Male, day 2 | *t*=5.011, *df*=8 | ** | 0.0010 |
|  | Female, day 3 | *t*=0.4795, *df*=9 | ns | 0.6430 |
|  | Male, day 3 | *t*=5.852, *df*=7 | *** | 0.0006 |
|  | Female, day 4 | *t*=0.7748, *df*=9 | ns | 0.4583 |
|  | Male, day 4 | *t*=3.116, *df*=7 | * | 0.0169 |
| Lateral<br>(L-AUC) | Comparison | *t*, *df* | Summary | *P* |
|  | Female, day 1 | *t*=0.4160, *df*=9 | ns | 0.6872 |
|  | Male, day 1 | *t*=1.975, *df*=9 | ns | 0.0797 |
|  | Female, day 2 | *t*=0.7012, *df*=9 | ns | 0.5009 |
|  | Male, day 2 | *t*=0.5277, *df*=8 | ns | 0.6120 |
|  | Female, day 3 | *t*=0.5298, *df*=9 | ns | 0.6091 |
|  | Male, day 3 | *t*=0.7421, *df*=7 | ns | 0.4822 |
|  | Female, day 4 | *t*=0.4336, *df*=9 | ns | 0.6748 |
|  | Male, day 4 | *t*=4.836, *df*=7 | ** | 0.0019 |
| Opposite<br>(O-AUC) | Comparison | *t*, *df* | Summary | *P* |
|  | Female, day 1 | *t*=0.6327, *df*=9 | ns | 0.5427 |
|  | Male, day 1 | *t*=2.133, *df*=9 | ns | 0.0617 |
|  | Female, day 2 | *t*=0.3549, *df*=9 | ns | 0.7308 |
|  | Male, day 2 | *t*=1.153, *df*=8 | ns | 0.2821 |
|  | Female, day 3 | *t*=0.3089, *df*=9 | ns | 0.7644 |
|  | Male, day 3 | *t*=1.405, *df*=7 | ns | 0.2028 |
|  | Female, day 4 | *t*=0.0202, *df*=9 | ns | 0.9843 |
|  | Male, day 4 | *t*=2.342, *df*=7 | ns | 0.0517 |

## Discussion

*App*^h/h^ rats of both sexes were able to adapt to the IntelliCage and acquire simple place learning and reversal tasks, as well as a more complex behavioral sequencing task, by 6-8 weeks of age. Males tended to perform better than females at 4-5 months of age in place learning with corner switch and behavioral sequencing. The results of the single corner restriction program for cohort B suggest that although individual variance exists among the rats, it is small enough that animals can be approximated as identical subjects for these IntelliCage experiments. Generating activity curves with aggregate cohort data is one way to reduce the impact of this variance on interpretation of cohort performance. Using AUC as a metric for comparing activity between groups is a natural extension of using linear fits on activity curves to estimate learning rate and takes full advantage of the data volume the IntelliCage offers. Task acquisition can be characterized by performance parameters—in this case, AUC—that are greater or less than the value that would be expected through chance alone, depending on the visit category. Chance C-AUC/O-AUC would be equal to the area of a right triangle with base of length 180 (number of minutes in a drinking session) and height of 0.25 (probability of visiting a correct/opposite corner at random), or 22.5. Similar calculations can be done for IC-AUC (180 × 0.75 = 67.5) and L-AUC (180 × 0.50 = 90). Significant session-wise differences in the appropriate direction can reflect task acquisition too, as seen with increases in C-AUC accompanied by decreases in IC-AUC, L-AUC, or O-AUC. These characteristics were observed for all the spatial learning programs except serial reversal, suggesting that the program is too complex for the rats to learn by 4-5 months of age. In general, a task that challenges the animals without being impossible to acquire would be ideal for identifying possible cognitive deficits in models of neurodegeneration and dementia. Task acquisition of behavioral sequencing but not serial reversal suggests that a program of intermediate difficulty using a sequence involving all four corners (in clockwise motion, for example) rather than just two in a single diagonal, might be worth testing in future studies. By these measures, this study establishes a baseline spatial learning profile for *App*^h/h^ control rats while providing initial validation of analytic methods exploring aggregate cohort activity and using AUC as a metric for task performance in the IntelliCage.

